# Genome-based targeted sequencing as a reproducible microbial community profiling assay

**DOI:** 10.1101/2020.08.07.241950

**Authors:** Jacquelynn Benjamino, Benjamin Leopold, Daniel Phillips, Mark D. Adams

## Abstract

Current sequencing-based methods for profiling microbial communities rely on marker gene (e.g. 16S rRNA) or metagenome shotgun sequencing (mWGS) analysis. We present a new approach based on highly multiplexed oligonucleotide probes designed from reference genomes in a pooled primer-extension reaction during library construction to derive relative abundance data. This approach, termed MA-GenTA: Microbial Abundances from Genome Tagged Analysis, enables quantitative, straightforward, cost-effective microbiome profiling that combines desirable features of both 16S rRNA and mWGS strategies. To test the utility of the MA-GenTA assay, probes were designed for 830 genome sequences representing bacteria present in mouse stool specimens. Comparison of the MA-GenTA data with mWGS data demonstrated excellent correlation down to 0.01% relative abundance and a similar number of organisms detected per sample. Despite the incompleteness of the reference database, NMDS clustering based on the Bray-Curtis dissimilarity metric of sample groups was consistent between MA-GenTA, mWGS and 16S rRNA datasets. MA-GenTA represents a potentially useful new method for microbiome community profiling based on reference genomes.

---

The primary molecular methods for determining microbial composition are based on marker gene sequencing or whole metagenome shotgun sequencing (mWGS). The 16S ribosomal RNA (rRNA) marker gene has been widely used for bacterial profiling for decades across diverse ecosystems^1,2^. Using this method, taxonomic classification of the bacterial community can be obtained at modest cost and a resolution that ranges from sub-species to family level, depending on the 16S rRNA segment that is sequenced^3–6^. Continued reduction in the cost of DNA sequencing has meant that mWGS approaches have become increasingly common due to the greater information on gene content, taxonomic resolution, and strain-level variation^7^, despite higher cost and complexity of data analysis.

The Human Microbiome Project^8^ and similar large-scale investments^9^ established methods and reference datasets for characterization of microbial profiles across diverse human body sites. As a result, the tools and reference genome datasets for characterizing human microbiomes are much better developed than for those involving other organisms. The mouse is widely used in microbiome studies that seek to demonstrate a causal role of microbes affecting a given trait and to understand the mechanisms by which microbes contribute to phenotypes^10^. The vast majority of mWGS sequences from mouse gut samples have no matches to named organisms in public databases^11^, substantially limiting the informativeness of this approach.

One approach to the limited reference genome sequences is construction of *in silico* genomes based on computational sequence assembly of large mWGS datasets to create “metagenome assembled genomes” or MAGs^12–14^. The integrated Mouse Gut Metagenomic Catalog (iMGMC)^15^ is one such effort. Combining 1.3 Tbp of data from 298 mouse metagenomic libraries, Lesker, *et al.* assembled 1.2 million contigs; a subset of these could be grouped into 830 high quality MAGs (hqMAGs) that are predicted to be >90% complete and <5% contaminated based on the representation of single copy genes^16^.

Here we describe a new approach to metagenome profiling termed MA-GenTA (Microbial Abundances from Genome Tagged Analysis) that combines the specificity of mWGS analysis with a simplified laboratory and analytical workflow (Figure 1). The availability of custom-designed highly multiplexed pools of oligonucleotides (“oligos”) has opened possibilities for a range of new assay methods to specifically target microbes at the species, strain, and even gene level. We adapted the Allegro Targeted Genotyping assay’s single primer enrichment technology that is widely used for genotyping^17,18^ and implemented it as a quantitative, straightforward, and cost-effective method for profiling mouse microbial communities based on the iMGMC hqMAGs.

**Figure 1.**
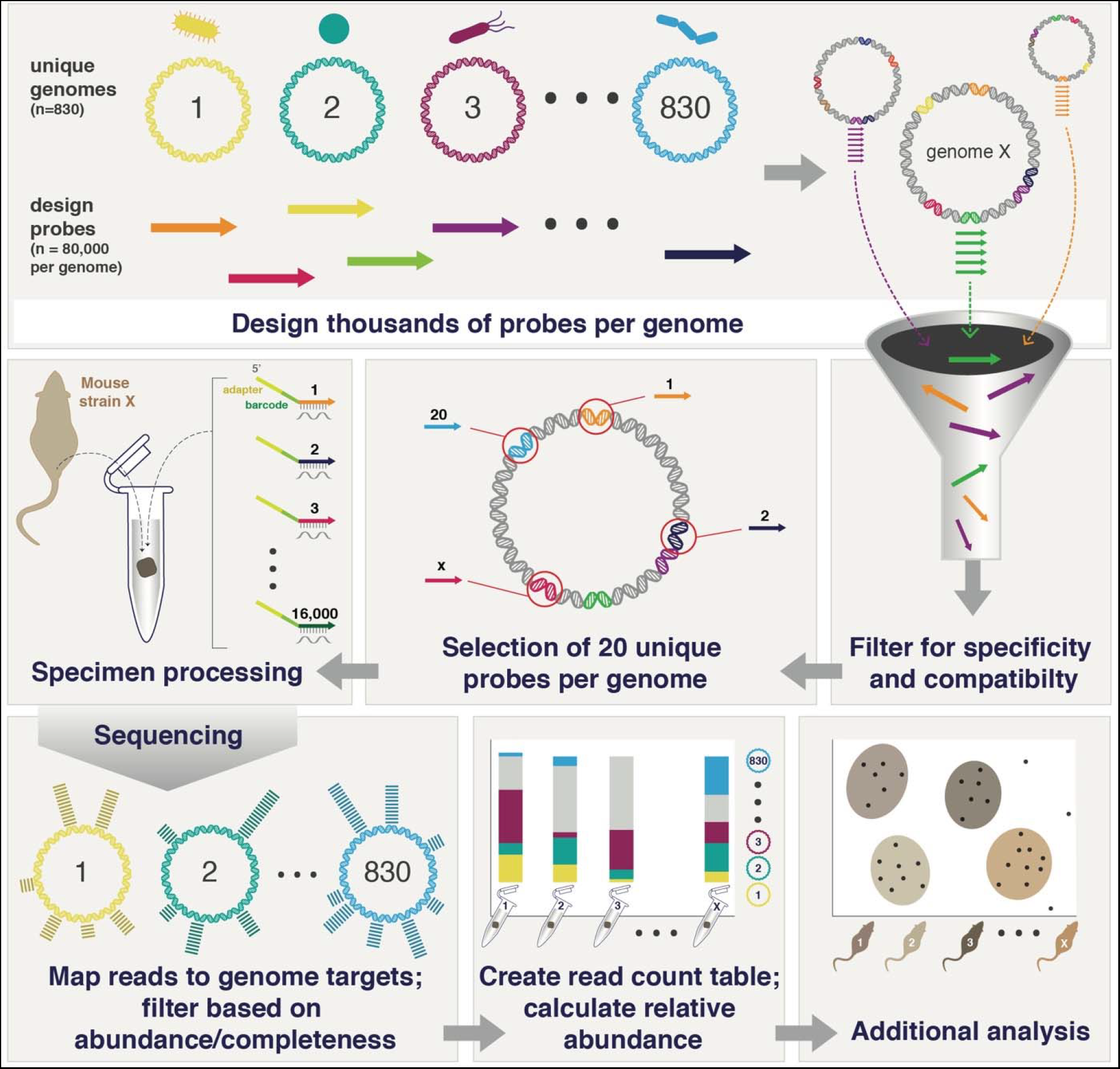
Overview of the MA-GenTA strategy. MA-GenTA utilizes software (CATCH) to design thousands of probes per genome for multiple genomes (830 in this study). All probes from the initial design are filtered based on multiple parameters (%GC, BLAST matches to inclusion/exclusion lists, non-unique matches across genomes, etc). Unique probes are selected for each genome (20 in this study). Probe pools are synthesized and used to prepare sequencing libraries using the Allegro Targeted Genotyping kit, and then sequenced. Reads are then mapped to the reference genomes to produce count tables for downstream analysis.

## Results

The MA-GenTA assay is based on approximating the relative abundance of hundreds of microbial species using sets of probes designed to be unique to each genome. The approach includes design of compatible probes directed at the genomes (or genes) of interest, library construction that uses the probe pools in a primer extension reaction, and integration of data across multiple probes to determine species abundance (Fig. 1). Oligonucleotide probe sets were designed using 830 iMGMC hqMAGs^15^. Preliminary results using a padlock probe design^19,20^ suggested that 20 probes per genome were sufficient to provide quantitative relative abundance information (data not shown). The padlock probe assay does not allow decoding of any additional adjacent sequence data for confirmation of probe specificity. We therefore sought to develop a method based on a single-primer extension assay, in which sequence adjacent to each probe is determined, allowing confirmation that the probe did in fact bind to the intended target.

Computational analysis suggests that each hqMAG is consistent with representing a single bacterial species and about 12% of hqMAGs are concordant with genome sequences of bacterial isolates that are present in GenBank. Most, though do not correspond with isolated bacteria, so in considering a probe design strategy, we decided to develop two completely independent probe sets for each hqMAG. We reasoned that concordance of relative abundance between these probe sets would provide additional support for the conjecture that the hqMAGs are reasonable approximations of *bona fide* genome sequences and that the organisms they represent are commonly found in the mouse gut.

Two defined-composition genomic DNA positive controls and a no-template negative control (NTC) were initially used to assess the specificity of each probe set. *Escherichia coli* gDNA and the ZymoBIOMICS Microbial Community Standard (Mock), which contains three species present in the iMGMC hqMAG set, one of which is an *E. coli* strain, were used as the positive controls.

Alignment of primary sequence reads showed that probes from many MAGs were detected for the Allegro and JAX designs for *E. coli* (493, 751), and Mock (264, 315) samples (grey dots in Fig. 2a). The vast majority of the MAGs matched in the *E. coli* and Mock samples were represented by a small number of probes with low relative abundance. After applying a probe-abundance threshold of ≥0.001% (Supplementary Fig. 1), there was only 1 MAG represented by >10 probes for both the Allegro and JAX designs in the *E. coli* sample and 3 and 2 MAGs for the Allegro and JAX designs in the Mock sample as expected (colored dots in Fig. 2a). For the *E. coli* sample, 99.95% and 99.28% of reads mapped to the *E. coli* genome for the Allegro and JAX designs, respectively. For the Mock community sample, 99.92% and 98.36% of reads mapped to the three genomes present in the Allegro design and two in the JAX design, respectively.

**Figure 2.**
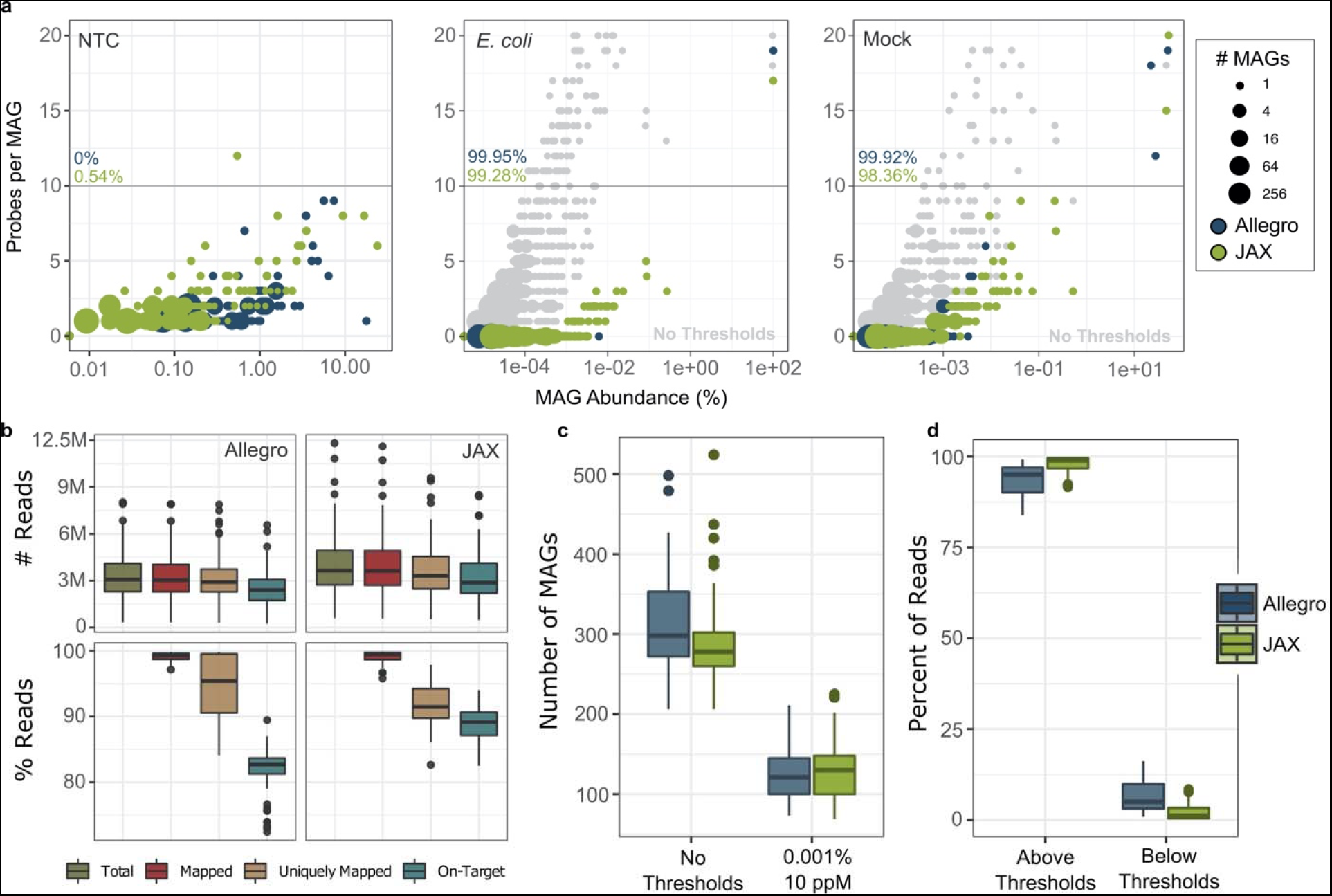
Use of control samples to establish thresholds for defining MAG presence. Thresholds for declaring a MAG present in a sample were determined using a no template control (NTC), *Escherichia coli* genomic DNA, and ZymoBIOMICS Microbial Community Standard. a, The number of probes present for each MAG (y-axis) and the MAG abundance (x-axis) for each control sample before applied thresholds is shown in gray. Blue (Allegro) and green (JAX) points indicate MAGs detected in each control sample after a 0.001% minimum probe-abundance threshold was applied. **b,** Sequencing reads from the Allegro and JAX probe pools were mapped to the iMGMC hqMAGs. **Top:** Read counts per sample for total reads, aligned reads, uniquely mapped reads, and uniquely-mapped, on-target reads. **Bottom:** Same data as in the top panel, but expressed as percent of total reads. **c,** The number of MAGs detected with minimum probe abundance and probe representation (probes per MAG-ppM) thresholds is shown compared to the number of MAGs detected with no thresholds across mouse samples. **d,** Most reads correspond to probes that pass the probe-representation thresholds.

In negative control samples, only a few thousand reads were obtained. NTC reads corresponded to 179 and 312 different probes and 77 and 138 MAGs in the Allegro and JAX designs, respectively (Fig. 2a). Of these probes, 94 (Allegro) and 142 (JAX) from *E. coli* overlapped with the NTC probes and 66 (Allegro) and 96 (JAX) from the Mock overlapped with the probes in the NTC. There are several potential sources of these reads: 1) contamination of the NTC with mouse stool DNA that was processed on the same batch; 2) contamination of the reagents used for library preparation; 3) self-annealing of primers within the probe set; or 4) sequencing-associated barcode-hopping. While there were many MAGs detected in the NTC, most of those MAGs were represented by only a few probes. No MAGs in the Allegro design and only one MAG in the JAX design had more than 10 probes represented (Fig. 2a). The MAG detected in the JAX dataset (single-China_7-4_110307.52) is a Muribaculaceae and present at high abundance in the majority of mouse samples.

The Allegro and JAX probe sets have no sequence overlap, thus they represent two completely independent assays for relative abundance of hqMAGs in mouse specimens. High concordance in probe representation and relative abundance would therefore support both the reliability of the MA-GenTA assay and the structural validity of the detected MAGs as representing a species present in the test sample. The Allegro and JAX probe sets were used to assay 72 mouse stool pellet samples, averaging 3.7 million sequencing reads per sample (Table 1, Supplementary Table 1). All reads for both datasets were mapped to the iMGMC hqMAGs reference. After mapping, reads that mapped to multiple regions were removed to produce uniquely mapped reads. The uniquely mapped reads were then filtered to include only reads that aligned adjacent to the designed probe region; this allowed us to determine probe-derived (on-target) reads. The two probe sets yielded similar numbers of sequencing reads and mapped reads (Fig. 2b). There was a larger variation in the proportion of uniquely mapped reads and fewer on-target reads in the Allegro dataset compared to the JAX dataset, suggesting that the JAX design pipeline may be more effective in selecting unique regions of each MAG. The previously chosen 0.001% minimum probe-abundance and 10 probes per MAG (ppM) thresholds were applied to the mouse samples (Fig. 2c). The number of MAGs observed in the mouse samples after applying the thresholds decreased by ~50% (Fig. 2d). However, over 90% of the reads matched MAGs present above the thresholds (Fig. 2d).

**Table 1.**
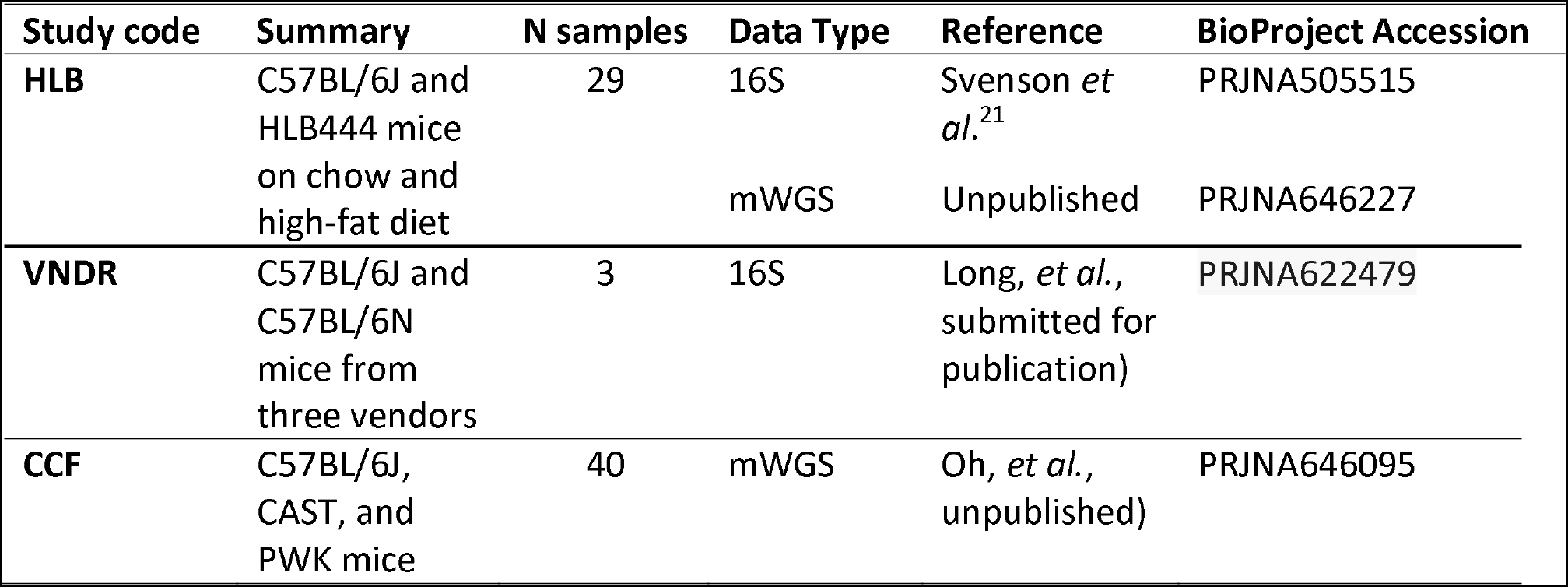
Mouse specimen groups used for analysis.

| Study code | Summary | N samples | Data Type | Reference | BioProject Accession |
| --- | --- | --- | --- | --- | --- |
| HLB | C57BL/6J and HLB444 mice on chow and high-fat diet | 29 | 16S | Svenson *et al.*<sup>21</sup> | PRJNA505515 |
|  |  |  | mWGS | Unpublished | PRJNA646227 |
| VNDR | C57BL/6J and C57BL/6N mice from three vendors | 3 | 16S | Long, *et al.*, submitted for publication) | PRJNA622479 |
| CCF | C57BL/6J, CAST, and PWK mice | 40 | mWGS | Oh, *et al.*, unpublished) | PRJNA646095 |

Comparison of the MAG abundances between the two designs without a probe abundance threshold gave a Pearson correlation coefficient of 0.98, demonstrating that the MAG abundance as measured by the Allegro and JAX probe sets were highly consistent (Fig. 3a). The points on the plot are colored by the number of probes detected in each MAG in both probe sets, showing higher abundance and better concordance between the probe sets for MAGs with reads from 10 or more probes. The MAGs were also plotted based on the number of probes detected in each dataset across all mouse samples, illustrating that MAGs tend to have high or low probe representation in both probe sets (Fig. 3b).

**Figure 3.**
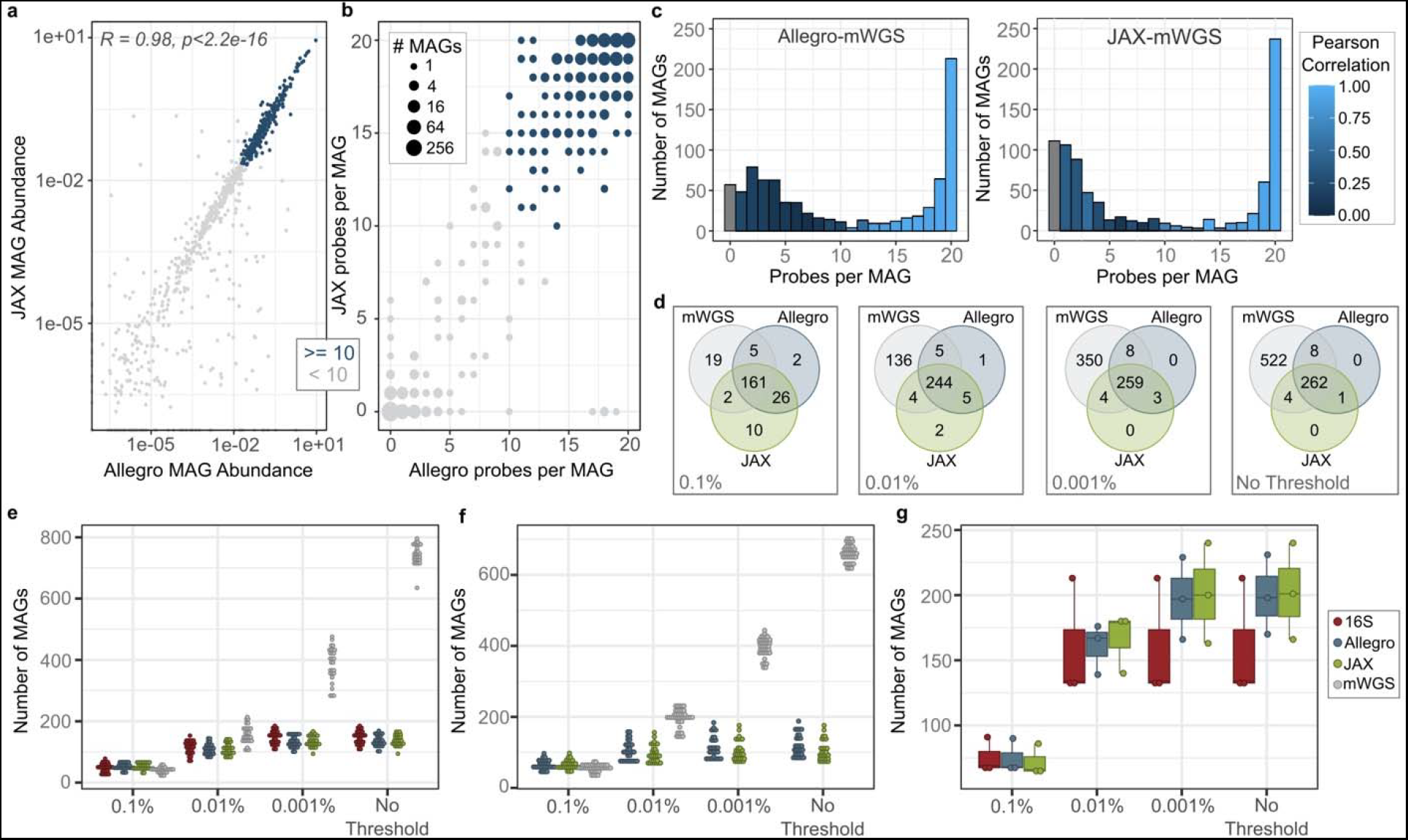
Comparison of MA-GenTA probe pools to established sequencing assays. **a,** The percent relative abundance of each MAG in each sample based on the Allegro design (x-axis) and the JAX design (y-axis) is shown. MAGs with 10 or more probes above the 0.001% probe-abundance threshold in both designs are shown in blue. Pearson correlation of the two designs is *R = 0.98*. **b,** The number of probes per MAG detected using the Allegro design (x-axis) and JAX design (y-axis) As in C, MAGs with at least 10 probes with ≥0.001% abundance in both assays are colored blue. Most MAGs have ≥15 probes per MAG above the threshold (top right) or ≤5 (bottom left). **c,** The relative abundance of each MAG as inferred from the targeted and mWGS data was compared across the mouse stool samples using histograms showing the number of MAGs (y-axis) with the number of probes observed per MAG (x-axis) with no minimum probe-abundance threshold. The color-scale shows the Pearson correlation of the relative abundance between the Allegro (left) JAX (right) data and the mWGS data. **d,** The total number of MAGs present in each assay (JAX, Allegro, mWGS) are shown in Venn-diagrams, highlighting the overlapping MAGs between the assays. **e,** Samples from the HLB dataset are shown with 16S rRNA v1-v3 OTUs, and hqMAGs detected by Allegro, JAX, and mWGS assays at a range of minimum probe-abundance thresholds. **f,** CCF samples with hqMAGs detected by Allegro, JAX, and mWGS assays. **g,** VNDR samples with 16S rRNA v1-v3 OTUs, and hqMAGs detected by Allegro and JAX assays.

### Comparison of the MA-GenTA assay to other microbial community profiling assays

mWGS data was available for 69 mouse fecal samples, enabling correlation of relative abundance data for each MAG between the two assays. MAGs were separated into groups based on the number of probes observed by MA-GenTA in each sample (e.g. from 1 to 20) and a Pearson correlation was performed on each group of MAGs between the MA-GenTA and mWGS abundance data (Fig. 3c and Supplementary Fig. 2, 3, Supplementary Table 2). For both the Allegro and JAX datasets, MAGs with ≥15 probes detected have relative abundance correlations of R ≥ 0.9 to the mWGS data. MAGs represented by less than 10 probes had poor Pearson correlations between the relative abundance of MA-GenTA and mWGS data (*R ≤ 0.23* for Allegro and *R ≤ 0.52* for JAX). Poor correlation of MAGs with fewer probes could be due to poor probe performance, improperly assembled MAGs, pan-genome differences between the MAG and the organisms present in our samples, sequencing depth disparities between the MA-GenTA assay and mWGS, or inflated abundance values in mWGS caused by read-mapping hotspots or conserved regions.

16S rRNA gene sequencing, mWGS, and the MA-GenTA assay are distinct ways of determining the number of bacterial species present in a sample. We compared the number of observed MAGs from the MA-GenTA assay with the number of 16S rRNA v1-v3 OTUs and MAGs detected in the mWGS data across the mouse samples from three studies (Fig. 3d-g). A MAG was considered present if at least 10 probes had >0.001% probe abundance. These thresholds were used in subsequent analyses of mouse stool datasets. The sensitivity to detect a MAG depends upon sequencing depth (more reads means it is more likely reads from a low-abundance genome will be detected) and probe representation (if a MAG truly represents the genome of a species present in the sample, then reads from a large fraction of probes should be observed).

All the datasets were filtered with MAG/OTU relative abundance thresholds of 0.1%, 0.01%, 0.001%, and no threshold. The total number of MAGs across the all HLB samples was compared between the MA-GenTA (JAX and Allegro) assay and mWGS at each threshold (Fig. 3d). There was a steep increase in the number of mWGS MAGs as thresholds were lowered, while the MAGs in the JAX and Allegro assays increased slightly. The Venn diagram for each threshold shows high overlap of MAGs detected between JAX and Allegro MA-GenTA datasets, with an increasing number of low-abundance MAGs detected only in the mWGS assay. Within the HLB dataset, the Allegro and JAX MA-GenTA datasets yielded similar numbers of MAGs, which were also similar to the number of 16S OTUs across all thresholds on a per-sample basis (Fig. 3e). The mWGS data detected similar numbers of MAGs to the 16S and targeted data for the 0.1% and 0.01% relative abundance thresholds, but much larger numbers at the 0.001% cutoff and without an abundance threshold. This observation is consistent with data shown in Supplementary Fig. 4 where many MAGs had ≥ 0.01% relative abundance in the mWGS data (yellow tones), but lower abundance and <10 probes per MAG in both MA-GenTA datasets. The CCF dataset consisted of JAX, Allegro, and mWGS data (Fig. 3f). Similar patterns to the HLB data were seen, except that more MAGs were observed in the mWGS data than the MA-GenTA MAGs at a 0.01% threshold. Most CCF samples that had more MA-GenTA reads than mWGS reads; when the reference database was extended to include lower completeness MAGs, fewer hqMAGs were observed using mWGS reads, suggesting that non-specific mapping could explain some of the discrepancy (Supplementary Fig.5). In the VNDR dataset, only 16S rRNA data was available for comparison. For these samples, more MAGs were detected by the MA-GenTA than 16S OTUs at lower abundances (Fig. 3g).

In order to demonstrate the utility of the MA-GenTA assay in characterizing microbial profiles in an experimental context, we used the MA-GenTA datasets for analysis of the HLB samples. Prior results identified OTU differences between C57BL/6J mice and HLB444 mice, which carry a mutation in the *Klf15* gene, on both a standard chow diet and after introduction of a high-fat, high-sugar diet (HF)^21^. HLB444 mice are resistant to diet-induced obesity when fed the HF diet. To determine the ability of the MA-GenTA assay to differentiate these groups, the Bray-Curtis dissimilarity metric was applied to the 16S, mWGS, and MA-GenTA data of the same samples and viewed with non-metric multi-dimensional scaling (NMDS) plots (Fig. 4a). All assays showed samples clustered by diet (Chow vs. HF) and mouse strain (C57BL/6J vs. HLB444). PERMANOVA analysis for each of the sequencing assays confirmed significant clustering between mouse strain and diet: Allegro assay (*f = 2.6961, p = 0.0029*), JAX assay (*f = 13.629, p = 0.0009*), 16S (*f = 19.581, p = 0.0009*), mWGS (*f = 2.05, p = 0.0099*) (Supplementary Table 3).

**Figure 4.**
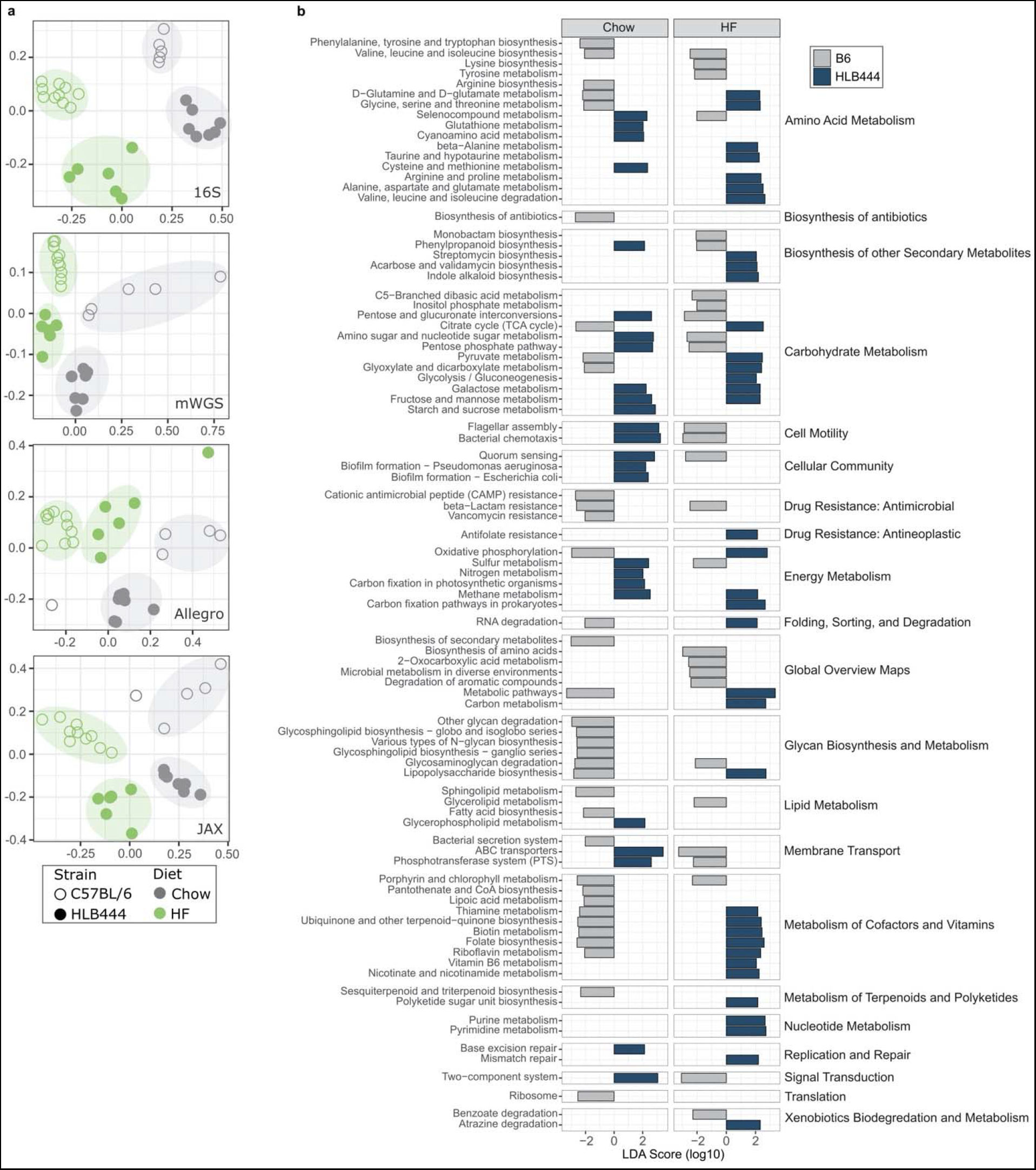
MA-GenTA as an assay for experimental group differentiation and functional analysis. **a,** The Bray-Curtis dissimilarity metric was applied to HLB data from each sequencing assay and shown in non-metric multi-dimensional scaling (NMDS) plots. Points are colored by diet, closed circles represent HLB444 samples, and open circles are C57BL/6J samples. All four sequencing assays cluster points based on diet and mouse strain. **b,** LDA analysis of KO pathways inferred by MA-GenTA MAG abundances shows differentially abundant pathways between HLB444 and B6 mouse strains on chow and HF diets.

### Functional analysis using MA-GenTA

Given the relative abundance of MAGs in each sample, we inferred the functional potential of each sample based on links of proteins encoded in each MAG to KEGG pathways. MA-GenTA read counts for each MAG in the HLB samples were assigned to KEGG pathways on a per-sample basis and then converted to relative abundance. Linear discriminant analysis in LEfSe was used to determine differentially abundant pathways between the two mouse strains and the two diets. The number of differentially abundant pathways varied across comparisons (HLB444 vs. B6 on HF diet (53,60), HLB444 vs. B6 on Chow (66,63), Chow vs. HF in HLB444 (101,103), and Chow vs. HF in B6 (75,81)) for the Allegro and JAX datasets respectively (Supplementary Table 4). Inter-assay KEGG pathway concordance was 82% for HLB444 vs. B6 on HF, 72% for HLB444 vs. B6 on Chow, 96% for Chow vs. HF in HLB444, and 77% for Chow vs. HF in B6. Consideration of the response of HLB444 and B6 strains to the HF diet showed differences in carbohydrate metabolism between the two strains on the HF diet, with HLB444 animals having higher representation of glycolysis, TCA cycle, and oxidative phosphorylation, and B6 animals with higher representation of pathways related to utilization of other sugars (Fig. 4b, Supplementary Figs. 6-13). These and other differences distinguished the response to HF diet of these two mouse strains and suggest microbial differences contribute to the ability of HLB mice to adapt to the HF diet.

### Specificity of MA-GenTA in a complex microbial environment

As an additional way to assess the specificity of probe targeting, both probe sets were used to assay metagenomic DNA extracted from a human stool specimen, which serves as a highly complex microbial sample with few organisms in common with mouse fecal bacteria (Supplementary Fig. 14). While there are deep-branching similarities in the gut microbiota of human and mouse, there are major differences at the genus and species level^11,22,23^. There were sixteen MAGs detected in the human stool sample using the same thresholds for detection as used for the mouse samples (minimum of 10 probes per MAG at ≥0.001% probe abundance). The taxa associated with the detected MAGs have previously been found in human stool samples^24–30^.

## Discussion

As the field of microbial community profiling grows, the need for informative, cost-effective, and streamlined assays of microbial composition becomes more important. Although initially developed for genotyping applications, we have shown that by combining results from multiple rigorously selected probes per genome, the Allegro Targeted Genotyping Assay can produce accurate microbial relative abundance data across at least three orders of magnitude dynamic range at a cost that is only moderately higher than 16S rRNA profiling. MA-GenTA bridges the gap between 16S rRNA gene sequencing and mWGS, combining some of the strengths of each approach (Table 2).

**Table 2.** Comparison of microbial community profiling assays.

| Feature | 16S rRNA gene sequencing | Whole metagenome sequencing | MA-GenTA |
| --- | --- | --- | --- |
| **Taxonomic Resolution** | ~Family/genus level for 16S rRNA subregions; strain level for full-length gene | Species/strain level | Species/strain level |
| **Gene content** | None | High | Inferred based on genome matches |
| **Analysis complexity** | Medium | High | Medium |
| **Cost** | <\$50/sample | >\$100/sample | \$50-\$75/sample |
| **Pros** | Quick community survey<br>Large number of studies from many environments/hosts | New organism/gene discovery<br>Direct comparison of datasets with same reference for mapping | Efficient pooled-sample workflow<br>Customized target selection/pool composition<br>Direct comparison of datasets with same reference for mapping |
| **Cons** | Limited taxonomic specificity<br>No gene content information | Possible mis-assignment of reads to closely related organisms<br>Cost | Limited to existing organisms/genomes<br>Limited pan-genome characterization |

A hallmark and major motivation of mWGS sequencing is the ability to analyze functional capability of the organisms in an environment. Strategies have been described to predict function based on OTU composition^31–33^, but they are strongly dependent on the reference databases and perform poorly on datasets from non-human-associated microbes^34^. Because probe design for the MA-GenTA assay requires reference genomes, this approach does not contribute to bacterial discovery. However, gene and pathway abundance data can be inferred from MA-GenTA data by pairing read counts to pathways represented in the reference genomes more directly than based on 16S rRNA sequences.

Capture-based targeted sequencing methods have been widely used for exome sequencing and cancer mutation profiling^17,18,35^, and represent a potential alternative approach for microbiome profiling. Guitor, *et al.* recently described a method for highly multiplexed detection of antibiotic resistance genes and bacteria that relies on biotinylated capture probes^36,37^. These probes and streptavidin bead capture kits are costly and require each specimen to be processed separately, making library preparation laborious. By contrast, the Allegro workflow involves pooling after a sample-specific tagging step and combination of pools can yield up to 3072 uniquely barcoded libraries on a single sequencing run. Up to 100k probes can be included in a single Allegro design. Unlike array-based platforms^38^, it is straightforward to alter the design of the MA-GenTA probe pool with each reagent order, allowing both the refinement of the selected probes for each genome and the inclusion of additional content over time.

The ability to synthesize probes based on user-defined parameters allows for broad or targeted study of microbial communities, specific species or strains, genes of interest, antibiotic resistance or virulence markers. Probe designs that focus on universal genes may be a good choice for species tagging, while probes targeting variable regions could provide additional information on pangenome variation. An important factor to consider when designing a probe pool for MA-GenTA is the reference database from which probes are chosen, including how representative the database is of organisms present in the sample. Across mouse mWGS samples, only about 60% of reads matched the iMGMC hqMAGs, reinforcing the need for a more robust reference for the mouse stool microbial community. Further optimization of the MA-GenTA assay might involve adjusting the number of probes per genome and how thresholds for probe abundance and probe representation are used to reduce noise and increase confidence of MAG assignment. Although not examined here, the specificity of the MA-GenTA assay would also be advantageous in specimens with high proportions of host genomic DNA where mWGS analysis is inefficient. The MA-GenTA assay could also be adapted to an RNAseq format for quantitative gene expression analysis.

## Methods

### Probe design and filtering

The “high quality” MAG set from the integrated Mouse Gut Metagenomic Catalog (iMGMC) was accessed from GitHub (https://github.com/tillrobin/iMGMC). The hqMAG set comprised 830 dereplicated genome equivalents predicted to be >90% complete and <5% contaminated based on analysis by CheckM^16^. Two probe design strategies were used. For the JAX design, the probe selection program CATCH^39^ was run on each hqMAG separately to design over 50,000 40-base probes per MAG. BLAST was used to match probes to Prokka-annotated ORFs^40^. Probes with BLAST matches shorter than 40 bp in length or less than 100% identity were removed, followed by probes corresponding to genome regions on a pre-defined discard list. Discard regions included annotations listed as tRNAs, ribosomal proteins, and with encoded proteins with the term “repeat” or “hypothetical” in the name. Probes were required to have between 45 and 65% G+C nucleotides. Probes with multiple matches within the hqMAG or to more than one hqMAG were also excluded. Probes matching the single-copy MUSiCC gene list^41^ were flagged for probe selection. All resulting probes were sent to Tecan Genomics (Redwood City, CA) where probe compatibility was assessed for probe pool production based on the Allegro Targeted Genotyping protocol, and probe pools with 20 probes per MAG were synthesized (JAX design), with 10 representing MUSiCC genes and 10 representing non-MUSiCC genes. The iMGMC hqMAGs were also used by Tecan Genomics to create a second probe pool (Allegro design) with 20 probes per MAG. There were 16 MAGs that did not pass probe-synthesis filtering metrics for the JAX design but were present in the Allegro design. The final probe pools contained 16,600 probes for the Allegro design and 16,280 probes for the JAX design. Cross-reference between the hqMAG set and the ZymoBIOMICs Microbial Community Standard was determined using BLAST alignment^42^, resulting in 3 MAGs matching genomes from the ZymoBIOMICS genomes (*Escherichia coli, Enterococcus faecalis, and Pseudomonas aeruginosa*).

### DNA Extraction of Mouse Stool Pellets and Controls

Genomic DNA isolated from mouse stool pellets from several studies was used for evaluation of the MA-GenTA assay (Table 2). All procedures used for animal husbandry and collection of specimens were approved by the Jackson Laboratory Animal Care and Use Committee and research was conducted in conformity with the *Public Health Service Policy on Humane Care and Use of Laboratory Animals*. The HLB and VNDR study pellets and positive controls (*E. coli*, ZymoBIOMICS Mock) were lysed using Qiagen PowerBead garnet tubes with 1 mL Qiagen InhibitEX buffer. The lysate was then processed with the QiaCube HT instrument using a modified Qiagen QIAamp 96 DNA QIAcube HT protocol^21^ (Svenson). Each sample (a single stool pellet, 10-60 mg total weight) was added to a Qiagen PowerBead 0.7 mm garnet tube with 1 mL of QIAGEN InhibitEX buffer. All samples were incubated at 65°C for 10 minutes followed by 95°C for 10 minutes. The samples were then mechanically lysed for 2 cycles of 30 seconds at 3,700 RPM on a QIAGEN Powerlyzer 24 Homogenizer, with a 1-minute rest period between cycles. Samples were then centrifuged at 10,000 × g for 1 minute, and then 200 μL of this lysate was then mixed with AL Buffer (285 μl) and Proteinase K (5 μL). The lysate was incubated for 10 minutes at 70°C and followed by an ice incubation for 5 minutes. 485 μL of lysate was transferred to a QiaCube HT instrument, where the lysate was combined with 200 μL of 100% Ethanol and then bound to the Qiamp 96 plate. Each well of the Qiamp 96 plate was then washed with 600 μL of AW1 Buffer, AW2 Buffer, and then 100% Ethanol. DNA was then eluted with 100 μL of AE Buffer without using TopElute fluid. The CCF stool pellets were homogenized with 500 μL Tissue and cell lysis buffer (Lucigen^©^) by pipetting up and down. An aliquot of 100 μL was removed and treated with an enzyme cocktail (5 μL 10 mg/mL lysozyme, 1 μL lysostaphin (5000 U/mL), 1 μL mutanolysin (5000 U/mL) and 20 μL Tissue and cell lysis buffer) for 30 minutes at 37°C. Buffer ASL (QIAGEN^©^) (200 μL with 0.5 μL anti-foaming agent DX) was added to each tube and mixed. Samples were placed on a QIAGEN^©^ TissueLyser II bead beater for 2× 3 minutes (30 Hz) and then spun down in a microcentrifuge. Each sample (200 μL) was further processed on the QIAGEN QIAamp 96 DNA QIAcube HT protocol.

### Allegro Targeted Genotyping Sample Prep and Sequencing

The Allegro Targeted Genotyping V2 protocol (publication number M01501, Tecan Genomics, Inc.) was followed for library preparation of all samples in duplicate with the Allegro and JAX probe pools. Briefly, gDNA samples were enzymatically fragmented, followed by ligation of barcoded adaptors. Barcoded samples were then purified and pooled together in groups of 48. Each pool of 48 samples was placed in an overnight annealing and extension reaction with the probe pool, followed by an AMPure XP bead purification. A qPCR step was used to determine the number of cycles used in the library amplification (18 cycles). Amplified libraries were bead purified (AMPure XP) and pooled in equimolar ratios for sequencing. A no template control (NTC), *Escherichia coli* gDNA (ATCC^®^ 8739™), a human stool metagenome DNA sample^43^ (Petersen et al), and a defined composition microbial community control (ZymoBIOMICS Microbial Community Standard, Cat # D6300) were used as controls. Libraries created from the Allegro Targeted Genotyping Assay were pooled and sequenced on an Illumina NovaSeq SP 2×150bp run, using the custom R1 primer and 1% spike-in of phiX174 library as recommended. Libraries were loaded on the NovaSeq SP at 60% of standard loading per Allegro Targeted Genotyping Assay recommendation; only forward read data was used for analysis.

### Data analysis

#### mWGS read mapping and 16S OTU generation

The raw mWGS sequences were trimmed of adapters and low-quality bases using Cutadapt version 1.14^44^. Host contaminant sequences were identified and filtered out using Kraken2 version 2.0.8-beta^45^. The clean sequences were aligned against the reference (iMGMC MAGs) using BWA version 0.7.12^46^ with parameter settings: bwa mem -M -P. The non-primary alignment reads were then filtered out using SAMtools version 0.1.19^47^ with parameter setting: -F 256. Reads were filtered using 97.5% ID and 50% coverage thresholds. Finally, the read count table by bin for each sample was generated from the alignment file. On average, about 60% of total mWGS reads mapped to the iMGMC 830 hqMAGs. 16S OTUs were generated for the HLB and VNDR data with USEARCH, using previously published parameters^21,48^.

#### MA-GenTA read mapping and data analysis

Raw sequences were trimmed using TrimGalore/CutAdapt to remove the 40 bp probe (https://github.com/FelixKrueger/TrimGalore)^44^. Read mapping to hqMAGs was performed using BWA. Sequences of up to 110 bp downstream of the probes were mapped to the iMGMC reference index. Reads mapped with <95.5% identity and ≤50% query length were removed. Secondary alignments with lower alignment scores were removed and then reads mapped to multiple sites with similar alignment scores were removed, which resulted in uniquely mapped reads. BEDtools intersect command was used to match read alignment locations to the genome locations of the designed probes to provide “on-target” read counts, removing reads that aligned to regions outside of the expected probe annealing location^49^. Counts tables were created representing the on-target read count and relative abundance of each probe in each hqMAG and the summed read counts and relative abundance for all probes per hqMAG. All analyses were performed in R (version 4.0.2)^50^. Allegro and JAX designs were compared based on the relative abundance per MAG and the number of probes per MAG matched in each sample. A Pearson correlation was performed on the MAG abundance comparison between the two designs and between each design and the relative abundance based on mWGS sequencing. The JAX and Allegro data were compared to 16S and mWGS data for the same samples on the basis of alpha (observed) and beta diversity (Bray-Curtis dissimilarity) metrics using Phyloseq^51^.

#### Functional analysis

Protein coding sequences in the hqMAGs were predicted using Prodigal^52^, implemented in Prokka^40^. Functional annotation of the predicted CDS regions was performed using EggNOG-Mapper^53^, using Diamond^54^ for searches, and with overlap parameters requiring at least 25% query and reference coverage. For each sample, the number of reads mapping to each MAG was assigned to each KEGG pathway^55^ for all constituent CDS regions. Differences in pathway abundance among sample groups was determined using linear discriminant analysis effect size with LEfSe^56^.

## Data Availability

Sequence data created in this study have been deposited in GenBank with the BioProject accession PRJNA646241. The probe sequences used for this study have been deposited to GitHub: https://github.com/TheJacksonLaboratory/MA-GenTA.

## Code Availability

All code used for probe design and data analysis, along with read count tables have been deposited to GitHub: https://github.com/TheJacksonLaboratory/MA-GenTA.

## Acknowledgements

We gratefully acknowledge the contribution of the Microbial Genomics Service and Genome Technologies Service at The Jackson Laboratory for expert assistance with the work described in this publication. We also gratefully acknowledge the Bioinformatics team at Tecan Genomics for their assistance in probe pool design and analysis development. We thank Julia Oh and John Graham for pre-publication access to mWGS data from stool specimens of collaborative cross founder (CCF) mouse strains.

