## Supplemental Figures for "Genome-based targeted sequencing as a reproducible microbial community profiling assay"

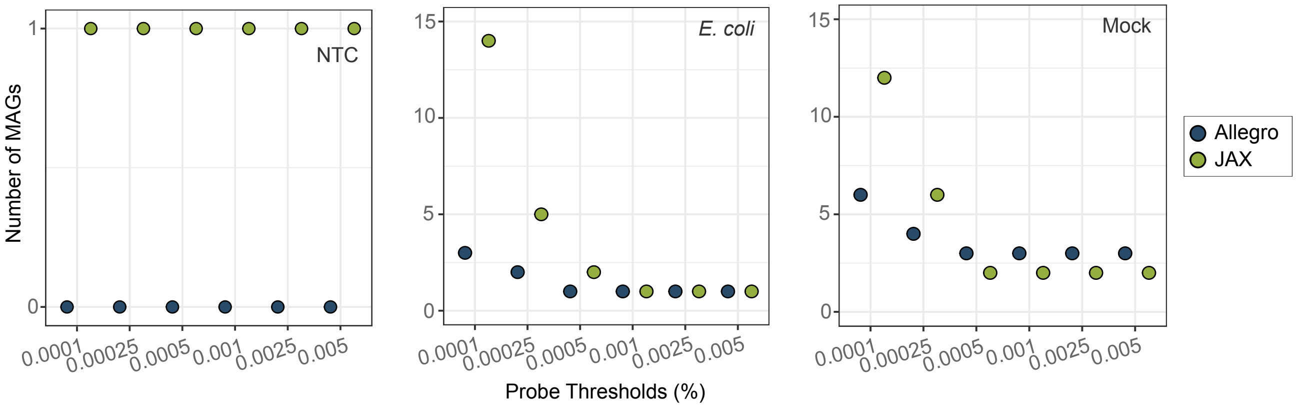


Supplementary Figure 1. Thresholds for declaring a MAG present in a sample were determined using a no template control (NTC), *Escherichia coli* genomic DNA, and ZymoBIOMICS Microbial Community Standard. Multiple probe-abundance thresholds were applied to the control samples (x-axis). The number of MAGs with ≥10 probes at each minimum probe-abundance threshold is shown on the y-axis. A probe abundance threshold of 0.001% improves accurate detection of MAGs.

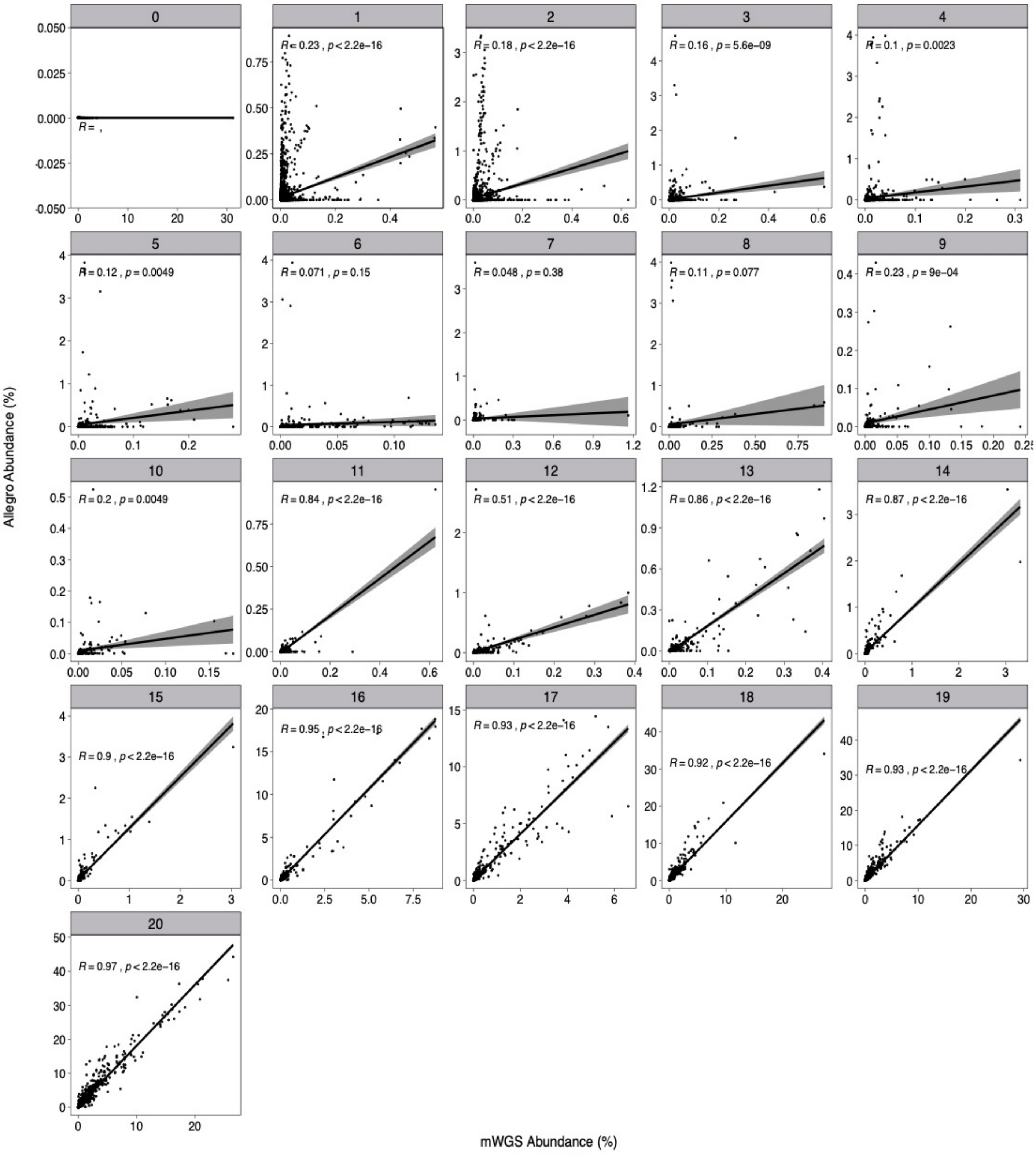


**Supplementary Figure 2. Correlation plots of the Allegro targeted data and mWGS data.** MAG abundance data was separated into groups based on the number of probes per MAG (plot titles in gray). The abundance of each MAG in the group was compared the MAG abundance determined by mWGS by Pearson Correlations. The correlation values for each group are at the top of each plot.

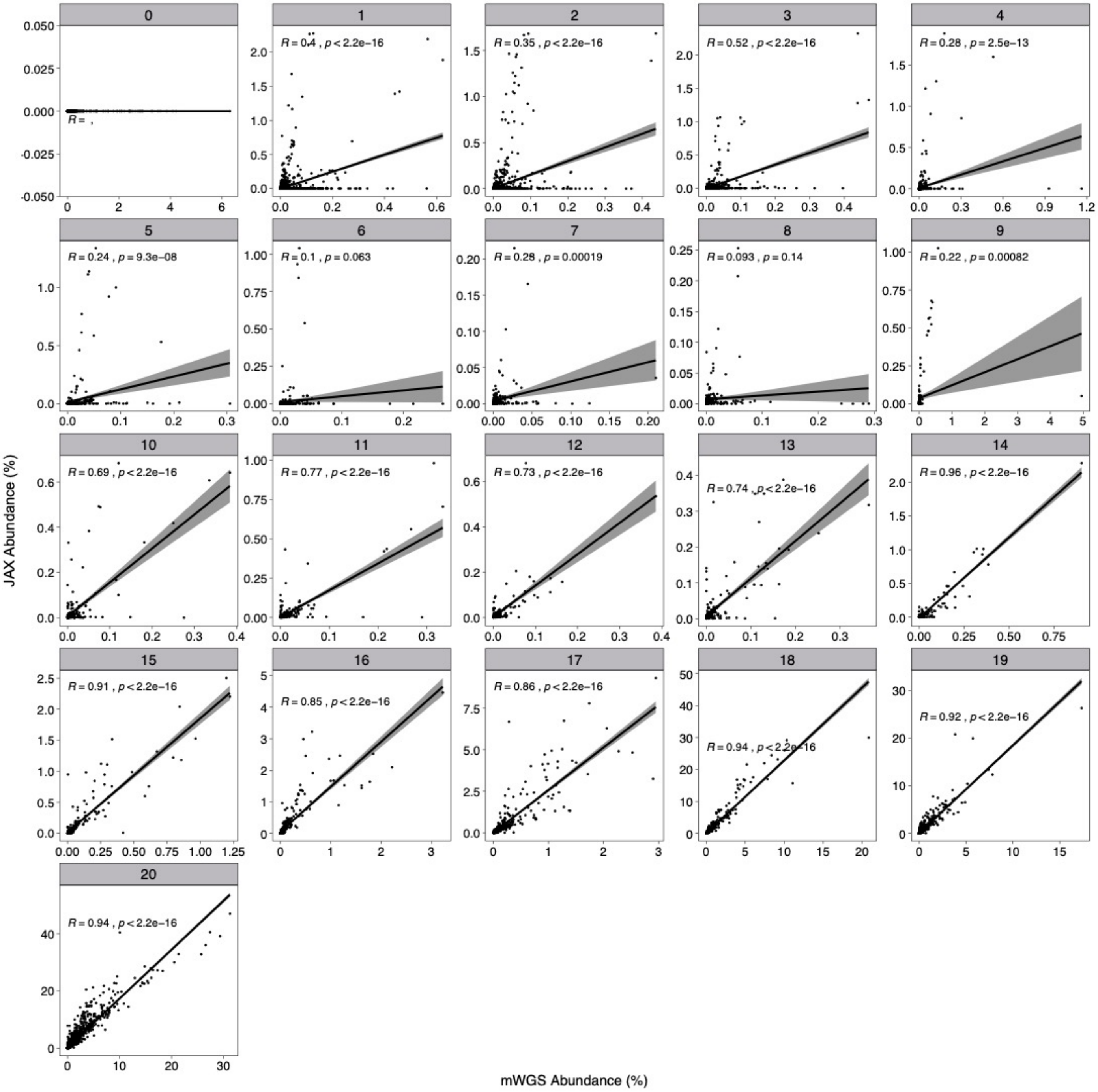


**Supplementary Figure 3. Correlation plots of the JAX targeted data and mWGS data.** MAG abundance data was separated into groups based on the number of probes per MAG (plot titles in gray). The abundance of each MAG in the group was compared the MAG abundance determined by mWGS data by Pearson Correlations. The correlation values for each group are at the top of each plot.

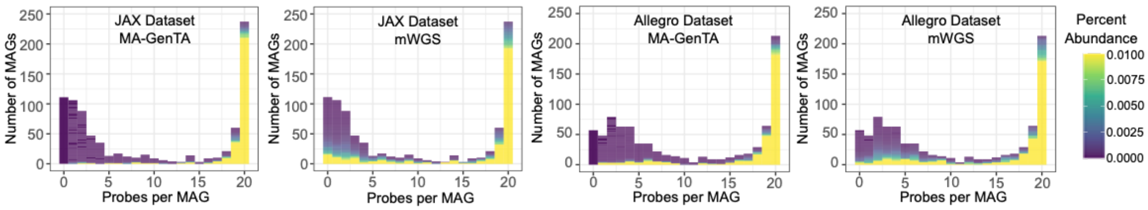


**Supplementary Figure 4. Comparison of MAG abundance between MA-GenTA and mWGS.** Number of MAGs (y-axis) were plotted against the number of probes per MAG (x-axis) for the Allegro and JAX data. Each MAG was colored by the percent abundance inferred by MA-GenTA and mWGS sequencing. Note that these graphs are the same as those shown in Figure 3c, but colored by relative abundance rather than Pearson correlation.

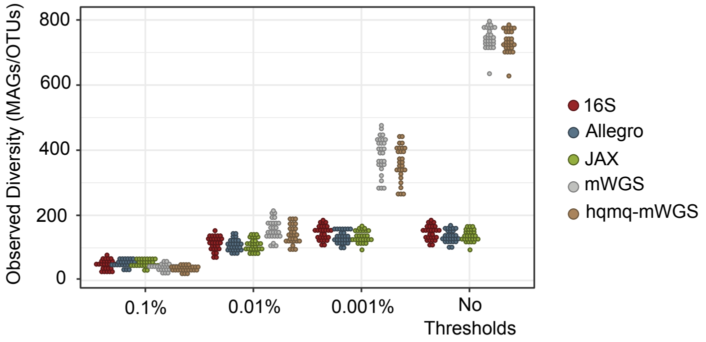


**Supplementary Figure 5. Observed diversity of HLB samples at various MAG/OTU abundance thresholds.** HLB samples sequenced by 16S, mWGS, and targeted sequencing assays were subjected to MAG/OTU abundance thresholds and the number of MAGs/OTUs are shown per sample on the y-axis. A ≥10 probes per MAG threshold was applied to the JAX and Allegro data before plotting. The mWGS data was mapped to the iMGMC high-quality (hq) MAGs (gray) and to the iMGMC hq+mq (high quality + medium quality) MAGs (brown). The observed diversity of the hqmq-mWGS samples shows only MAGs present in the probe design for the JAX and Allegro assay (iMGMC hq). The observed diversity of the hqmq-mWGS samples is lower due to reads mapping to MAGs not present in the iMGMC hq set.

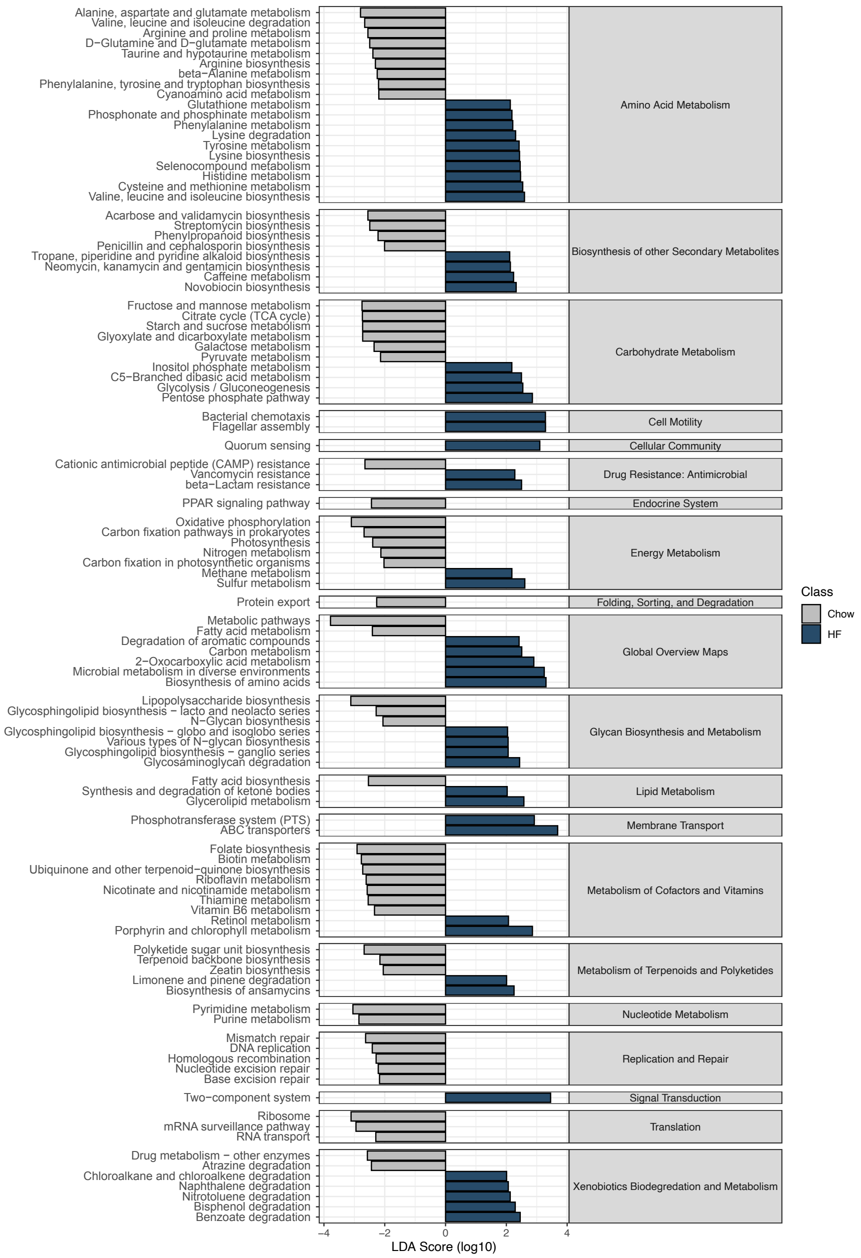


**Supplementary Figure 6. Differentially abundant pathways between chow and HF diets in HLB444 mice (JAX).** LDA analysis of KO pathways inferred by MA-GenTA MAG abundances shows differentially abundant pathways.

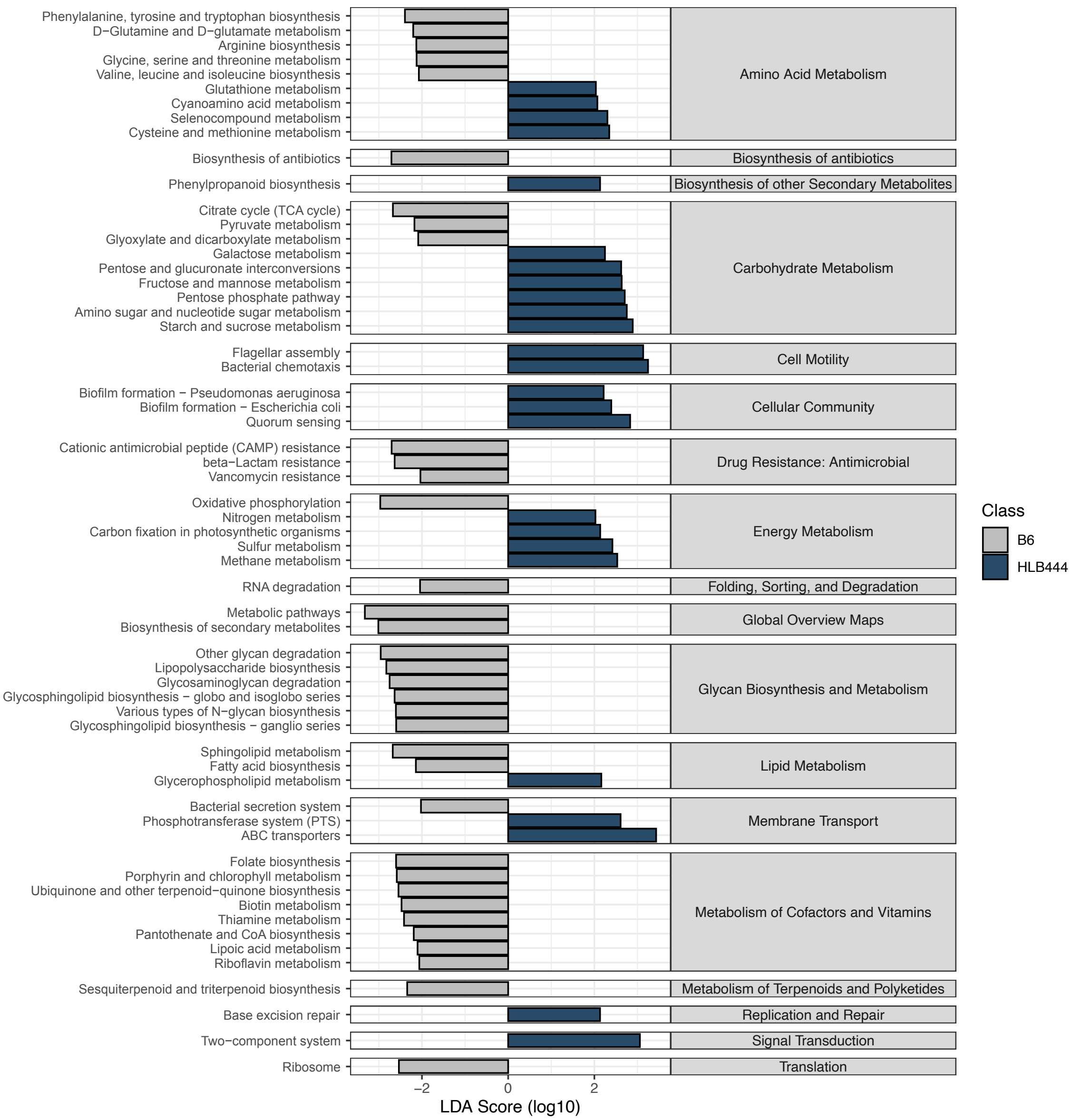


**Supplementary Figure 7. Differentially abundant pathways between HLB444 and C57BL/6J mice on a HF diet (JAX).** LDA analysis of KO pathways inferred by MA-GenTA MAG abundances shows differentially abundant pathways.

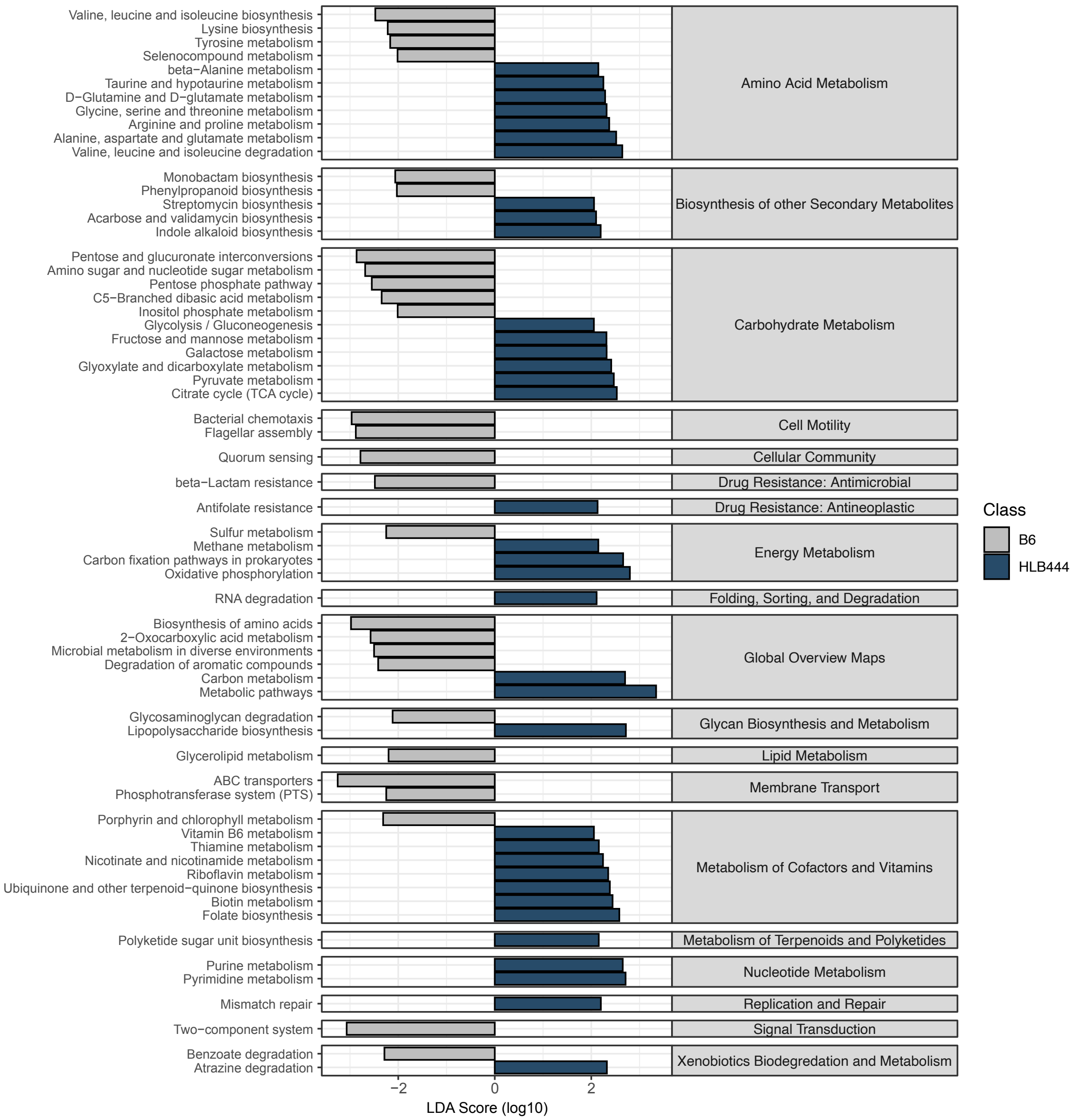


**Supplementary Figure 8. Differentially abundant pathways between HLB444 and C57BL/6J mice on a chow diet (JAX).** LDA analysis of KO pathways inferred by MA-GenTA MAG abundances shows differentially abundant pathways.

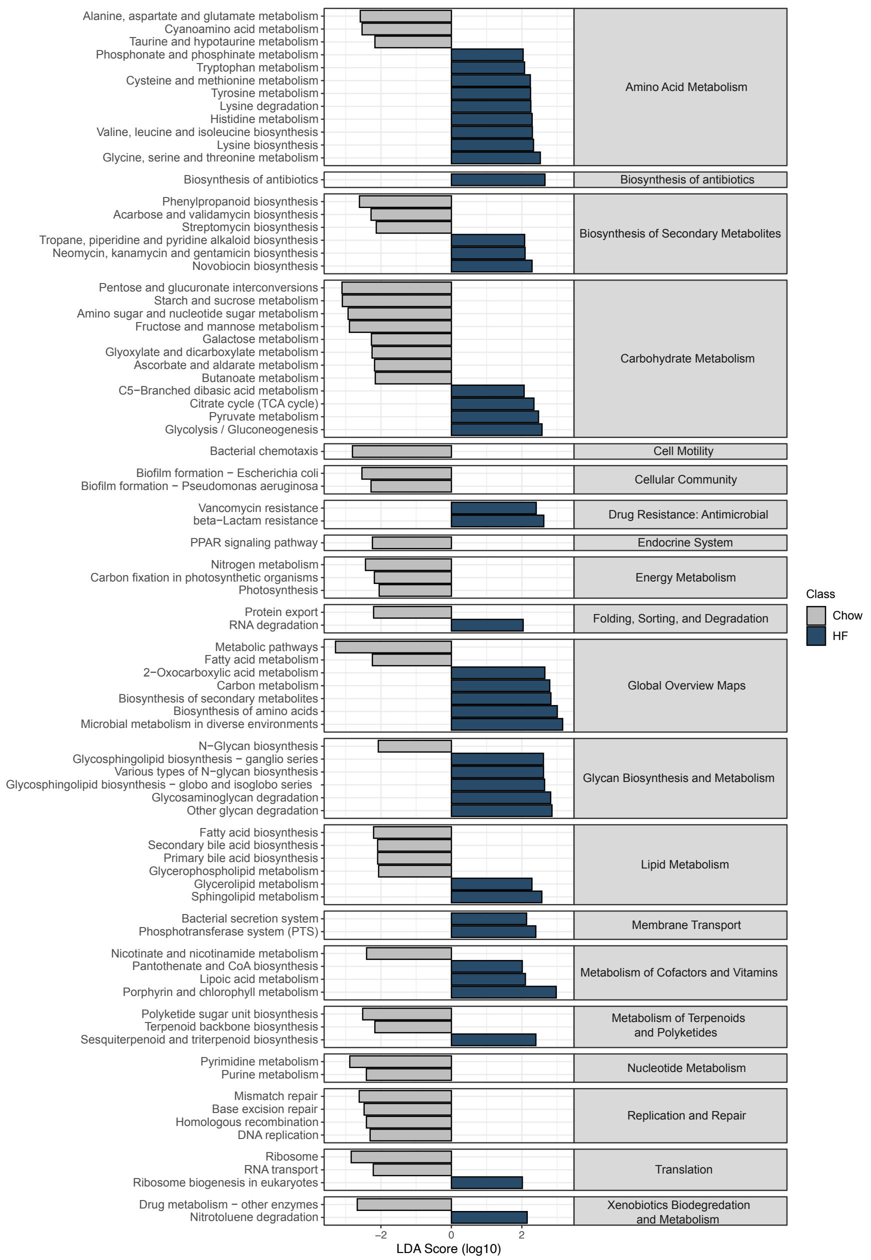


**Supplementary Figure 9. Differentially abundant pathways between chow and HF diets in C57BL/6J mice (JAX).** LDA analysis of KO pathways inferred by MA-GenTA MAG abundances shows differentially abundant pathways.

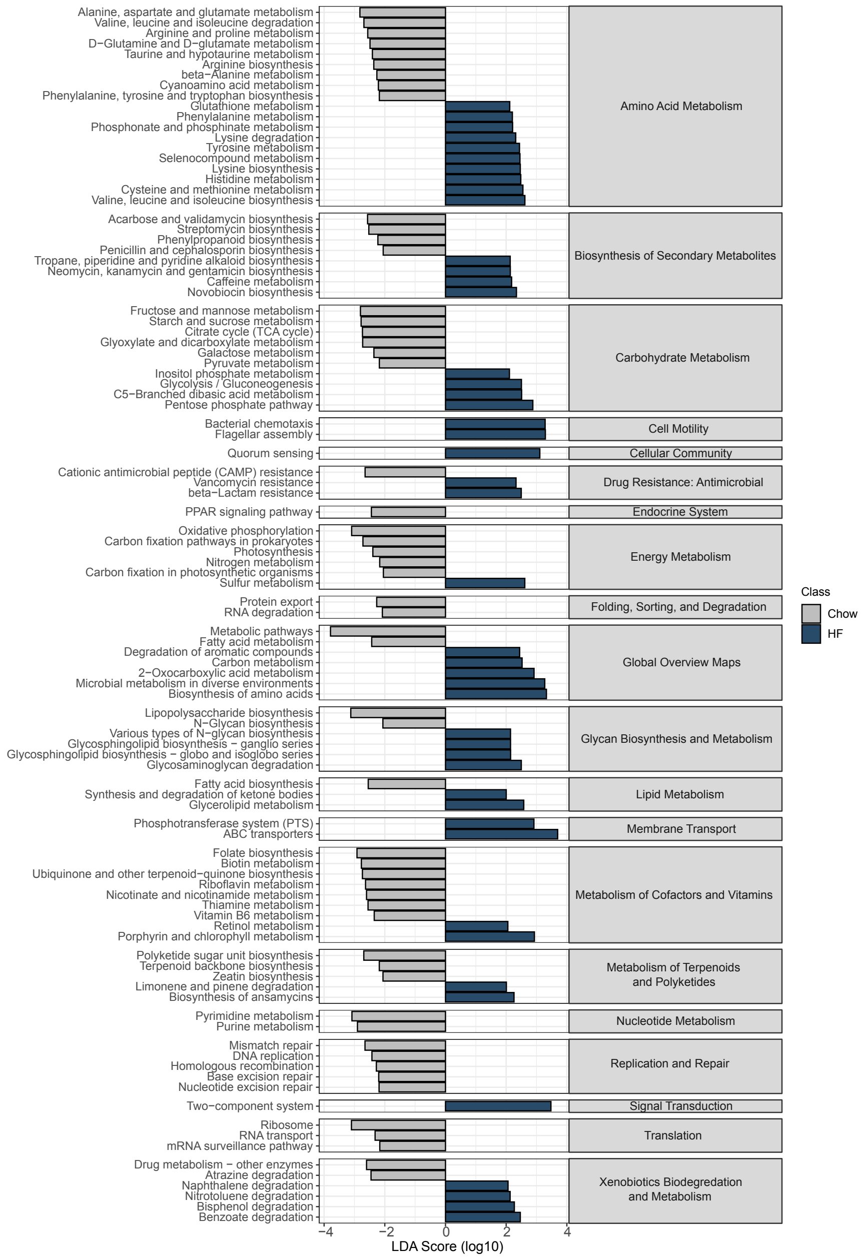


**Supplementary Figure 10. Differentially abundant pathways between Chow and HF diets in HLB444 mice (Allegro).** LDA analysis of KO pathways inferred by MA-GenTA MAG abundances shows differentially abundant pathways.

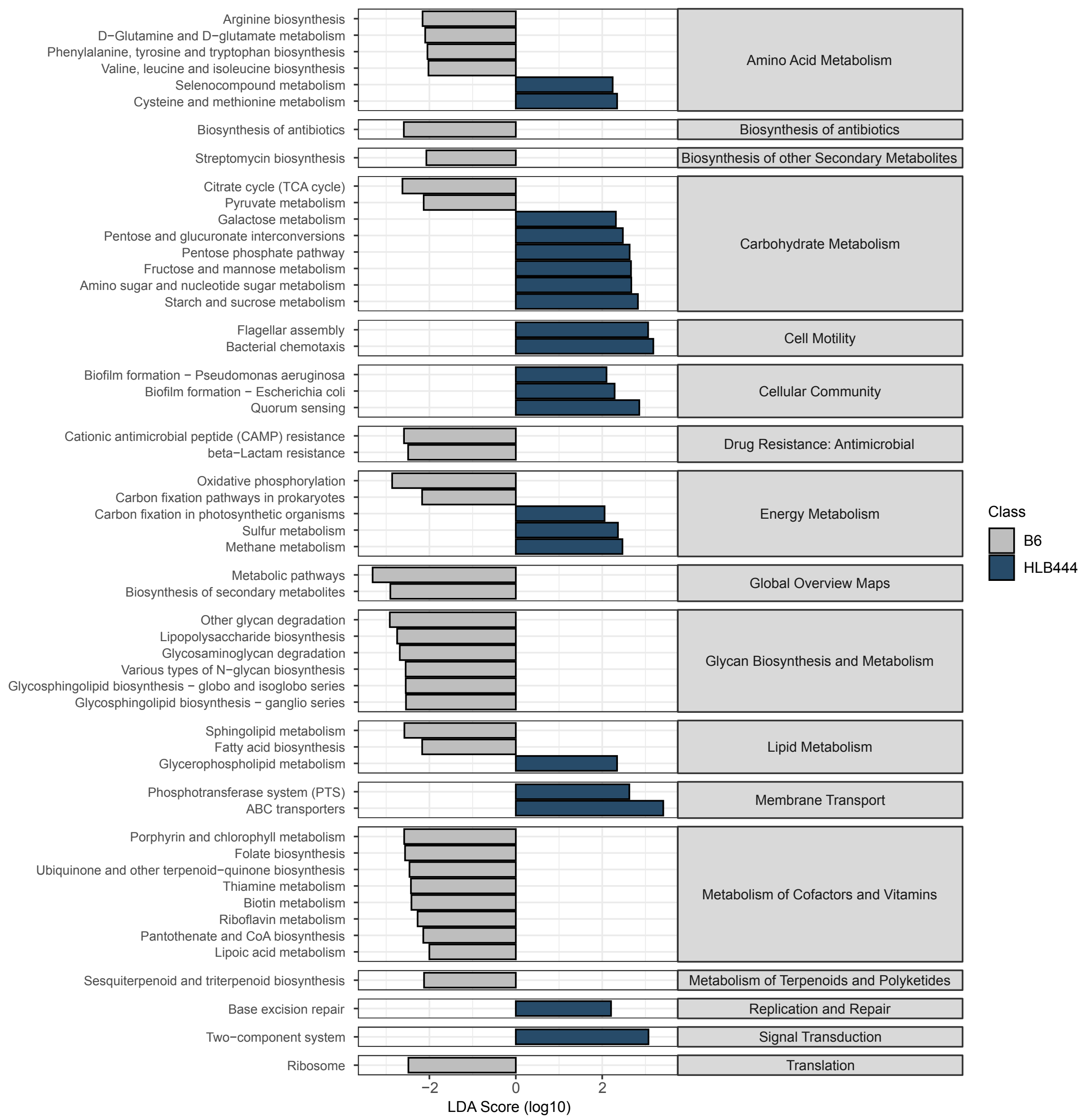


**Supplementary Figure 11. Differentially abundant pathways between HLB444 and C57BL/6J mice on a HF diet (Allegro).** LDA analysis of KO pathways inferred by MA-GenTA MAG abundances shows differentially abundant pathways.

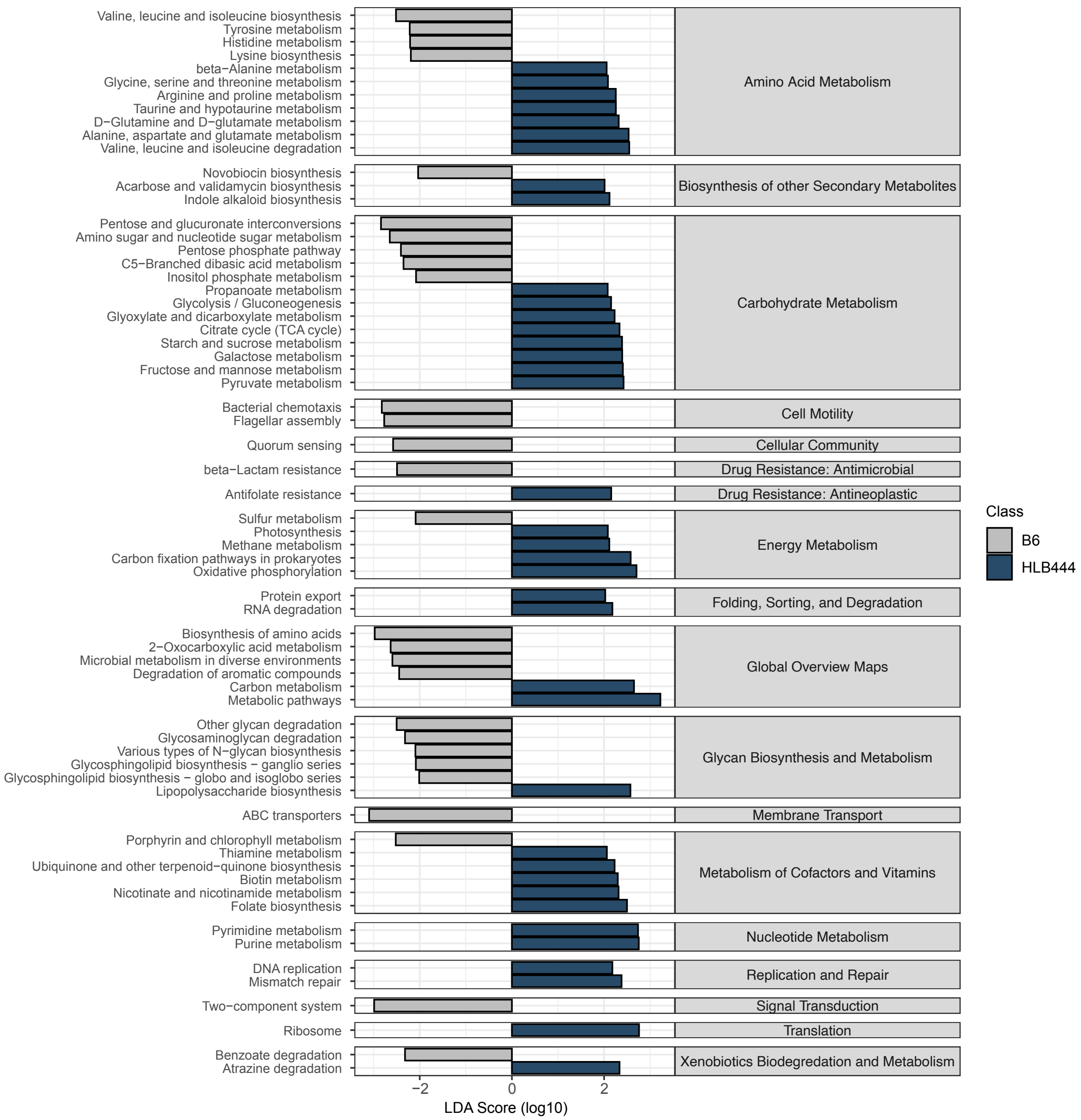


**Supplementary Figure 12. Differentially abundant pathways between HLB444 and C57BL/6J mice on a chow diet (Allegro).** LDA analysis of KO pathways inferred by MA-GenTA MAG abundances shows differentially abundant pathways.

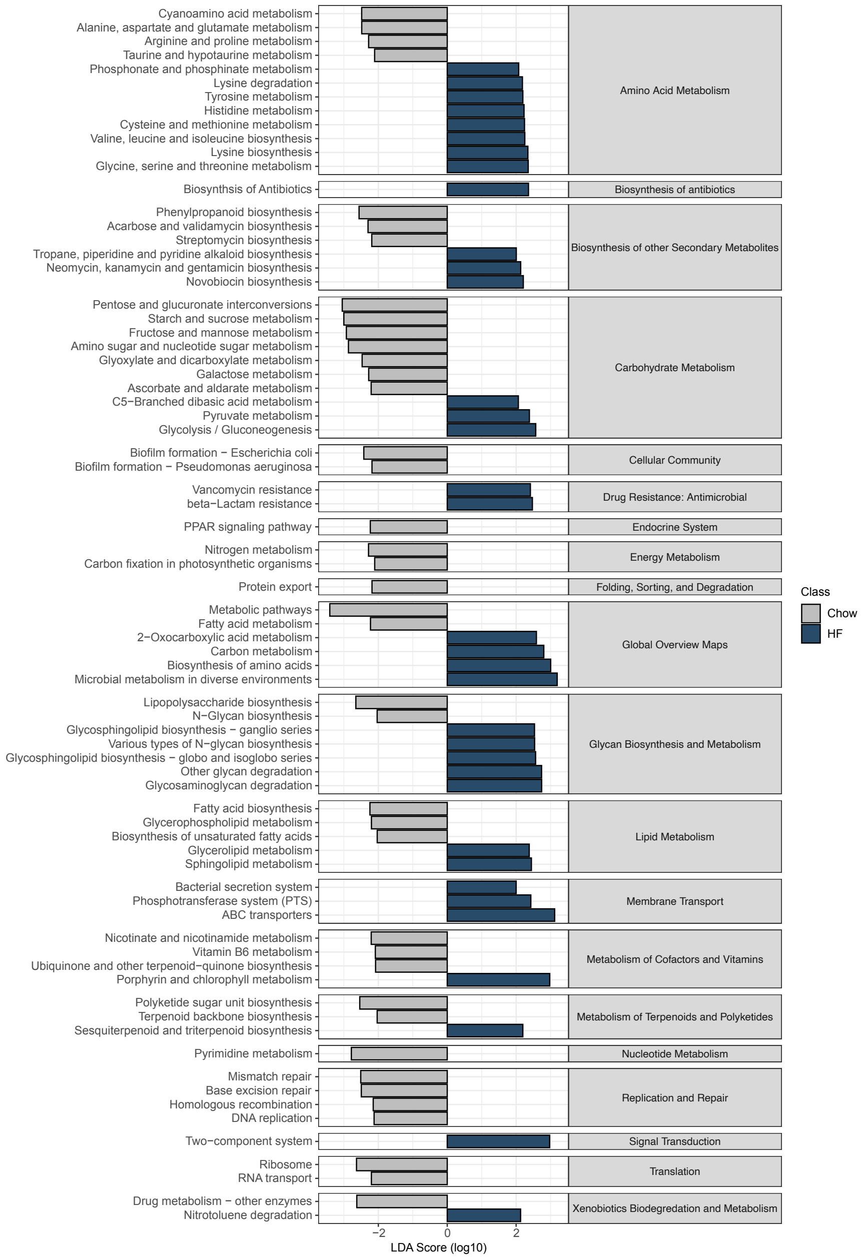


**Supplementary Figure 13. Differentially abundant pathways between Chow and HF diets in C57BL/6J mice (Allegro).** LDA analysis of KO pathways inferred by MA-GenTA MAG abundances shows differentially abundant pathways.

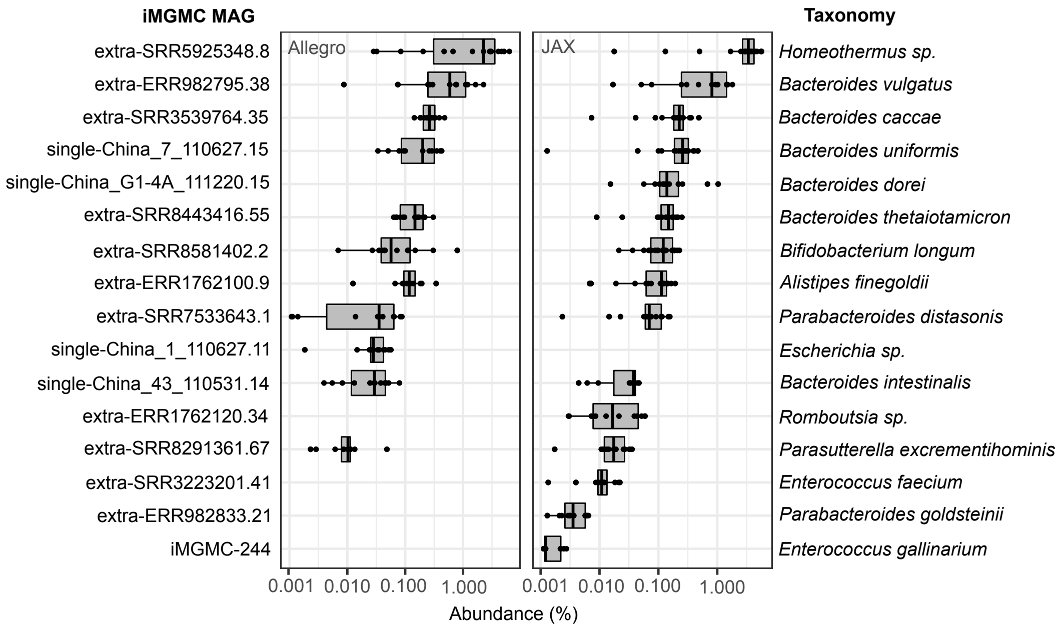


**Supplementary Figure 14. MA-GenTA as a precise assay.** The Allegro and JAX probe pools were used on a human sample to test the detection reliability of the probes in a complex sample. MAGs shown have at least 10 probes present at ≥0.001% relative abundance.

Supplementary Table 2. Pearson Correlation of MA-GenTA vs. mWGS.

| # Probes per MAG | Allegro | JAX |
|---|---|---|
| 1 | 0.23 | 0.4 |
| 2 | 0.18 | 0.35 |
| 3 | 0.16 | 0.52 |
| 4 | 0.1 | 0.28 |
| 5 | 0.12 | 0.24 |
| 6 | 0.071 | 0.1 |
| 7 | 0.048 | 0.28 |
| 8 | 0.11 | 0.093 |
| 9 | 0.23 | 0.22 |
| 10 | 0.2 | 0.69 |
| 11 | 0.84 | 0.77 |
| 12 | 0.51 | 0.73 |
| 13 | 0.86 | 0.74 |
| 14 | 0.87 | 0.96 |
| 15 | 0.9 | 0.91 |
| 16 | 0.95 | 0.85 |
| 17 | 0.93 | 0.86 |
| 18 | 0.92 | 0.94 |
| 19 | 0.93 | 0.92 |
| 20 | 0.97 | 0.94 |

Highlighted values: $p < 0.05$

Supplementary Table 3. PERMANOVA statistics of Bray-Curtis dissimilarity in different sequencing assays.

| Experimental Factors | df | SS | MS | Pseudo-F | $R^2$ | P |
|---|---|---|---|---|---|---|
| Allegro | | | | | | |
| Strain/Diet | 3 | 1.6571 | 0.5536 | 2.6961 | 0.2506 | 0.002997 |
| Residuals | 24 | 4.9169 | 0.20487 | | 0.74794 | |
| Total | 27 | 6.5740 | | | 1 | |
| JAX | | | | | | |
| Strain/Diet | 3 | 3.9962 | 1.33207 | 13.629 | 0.63012 | 0.000999 |
| Residuals | 24 | 2.3457 | 0.09774 | | 0.36988 | |
| Total | 27 | 6.3419 | | | 1 | |
| 16S | | | | | | |
| Strain/Diet | 3 | 4.1995 | 1.39984 | 19.581 | 0.70995 | 0.000999 |
| Residuals | 24 | 1.7157 | 0.007149 | | 0.29005 | |
| Total | 27 | 5.9152 | | | 1 | |
| mWGS | | | | | | |
| Strain/Diet | 3 | 1.1884 | 0.39613 | 2.0508 | 0.20405 | 0.00999 |
| Residuals | 24 | 4.6357 | 0.19316 | | 0.79595 | |
| Total | 27 | 5.8241 | | | 1 | |

PERMANOVA tests were performed using Strain and Diet as the group over 1000 permutations.
