## Supplemental Table 1 for "Genome-based targeted sequencing as a reproducible microbial community profiling assay"

| Sample | Total Reads | Mapped Reads | Unique Reads | On-Target Reads | Study Code | Additional Data | 16S Read Count | mWGS Mapped Reads |
|---|---|---|---|---|---|---|---|---|
| **CAST_L_T2_P1V2** | 3,538,249 | 3,463,218 | 3,008,160 | 2,477,332 | CCF | mWGS | | 1,199,883 |
| **CAST_L_T2_P1V4** | 4,128,629 | 4,075,711 | 3,620,097 | 3,024,607 | CCF | mWGS | | 1,199,883 |
| **B6J_2R1L_T2_P1V4** | 3,217,182 | 3,197,107 | 3,090,703 | 2,785,420 | CCF | mWGS | | 1,592,601 |
| **B6J_2R_T1_P2V2** | 3,238,300 | 3,215,252 | 3,194,358 | 2,691,469 | CCF | mWGS | | 1,263,735 |
| **B6J_2R_T1_P2V4** | 4,828,407 | 4,803,106 | 4,512,015 | 4,128,341 | CCF | mWGS | | 1,263,735 |
| **B6J_2R1L_T1_P2V2** | 3,315,661 | 3,295,397 | 3,279,764 | 2,646,630 | CCF | mWGS | | 2,180,489 |
| **B6J_2R1L_T1_P2V4** | 5,293,350 | 5,274,505 | 4,945,763 | 4,404,780 | CCF | mWGS | | 2,180,489 |
| **B6J_2R1L_T2_P1V2** | 2,743,736 | 2,710,214 | 2,601,317 | 2,263,225 | CCF | mWGS | | 1,592,601 |
| **B6J_3R_T1_P2V2** | 4,502,696 | 4,482,995 | 4,465,147 | 3,864,584 | CCF | mWGS | | 2,078,380 |
| **B6J_3R_T1_P2V4** | 5,413,147 | 5,394,310 | 5,196,451 | 4,751,589 | CCF | mWGS | | 2,078,380 |
| **B6J_3R_T5_P2V2** | 6,068,907 | 6,040,429 | 6,012,853 | 4,848,411 | CCF | mWGS | | 2,557,725 |
| **B6J_3R_T5_P2V4** | 4,799,462 | 4,779,150 | 4,547,947 | 4,093,635 | CCF | mWGS | | 2,557,725 |
| **B6J_B_T1_P2V2** | 2,654,000 | 2,601,851 | 2,554,927 | 2,108,249 | CCF | mWGS | | 1,118,157 |
| **B6J_B_T1_P2V4** | 2,517,711 | 2,474,516 | 2,267,359 | 2,065,729 | CCF | mWGS | | 1,118,157 |
| **B6J_L_T5_P2V2** | 4,754,565 | 4,738,546 | 4,712,663 | 3,866,190 | CCF | mWGS | | 2,077,001 |
| **B6J_L_T5_P2V4** | 6,121,807 | 6,101,981 | 5,967,043 | 5,473,721 | CCF | mWGS | | 2,077,001 |
| **B6J_R_T5_P2V2** | 2,489,055 | 2,478,987 | 2,470,029 | 2,024,308 | CCF | mWGS | | 1,659,624 |
| **B6J_R_T5_P2V4** | 2,536,981 | 2,527,998 | 2,442,305 | 2,258,105 | CCF | mWGS | | 1,659,624 |
| **CAST_2B_T1_P2V2** | 2,685,061 | 2,675,958 | 2,443,281 | 2,065,797 | CCF | mWGS | | 2,255,415 |
| **CAST_2B_T1_P2V4** | 2,262,321 | 2,256,367 | 2,103,224 | 1,894,700 | CCF | mWGS | | 2,255,415 |
| **CAST_2B_T5_P2V2** | 2,232,947 | 2,204,574 | 2,179,982 | 1,747,576 | CCF | mWGS | | 2,884,361 |
| **CAST_2B_T5_P2V4** | 2,612,646 | 2,577,040 | 2,356,541 | 2,100,283 | CCF | mWGS | | 2,884,361 |
| **CAST_2R1L_T1_P2V2** | 2,212,683 | 2,204,409 | 2,010,907 | 1,676,857 | CCF | mWGS | | 1,796,698 |
| **CAST_2R1L_T1_P2V4** | 2,276,619 | 2,269,419 | 2,106,040 | 1,883,781 | CCF | mWGS | | 1,796,698 |
| **CAST_2R1L_T2_P1V2** | 1,871,238 | 1,837,168 | 1,643,881 | 1,381,962 | CCF | mWGS | | 1,624,570 |
| **CAST_2R1L_T2_P1V4** | 2,643,994 | 2,594,733 | 2,328,648 | 2,030,119 | CCF | mWGS | | 1,624,570 |
| **CAST_2R1L_T5_P2V2** | 4,923,573 | 4,888,381 | 4,855,811 | 4,041,822 | CCF | mWGS | | 1,485,578 |
| **CAST_2R1L_T5_P2V4** | 4,967,176 | 4,938,501 | 4,773,656 | 4,387,351 | CCF | mWGS | | 1,485,578 |
| **CAST_3R_T5_P2V2** | 4,483,502 | 4,469,945 | 4,456,047 | 3,710,694 | CCF | mWGS | | 1,963,198 |
| **CAST_3R_T5_P2V4** | 4,933,032 | 4,920,595 | 4,756,506 | 4,355,956 | CCF | mWGS | | 1,963,198 |
| **CAST_B_T2_P1V2** | 3,029,171 | 3,007,803 | 2,629,445 | 2,162,203 | CCF | mWGS | | 1,698,096 |
| **CAST_B_T2_P1V4** | 3,920,378 | 3,901,545 | 3,477,083 | 2,880,738 | CCF | mWGS | | 1,698,096 |
| **CAST_B_T5_P2V2** | 2,742,127 | 2,730,979 | 2,720,707 | 2,227,446 | CCF | mWGS | | 1,556,930 |
| **CAST_B_T5_P2V4** | 3,297,866 | 3,286,678 | 3,161,407 | 2,874,609 | CCF | mWGS | | 1,556,930 |
| **CAST_L_T1_P2V2** | 2,572,514 | 2,552,764 | 2,406,680 | 2,033,586 | CCF | mWGS | | 1,934,741 |
| **CAST_L_T1_P2V4** | 3,655,883 | 3,635,047 | 3,414,573 | 3,168,668 | CCF | mWGS | | 1,934,741 |
| **CAST_N_T2_P1V2** | 1,643,600 | 1,624,265 | 1,454,447 | 1,196,743 | CCF | mWGS | | 1,355,359 |
| **CAST_N_T2_P1V4** | 1,625,977 | 1,614,315 | 1,446,293 | 1,229,234 | CCF | mWGS | | 1,355,359 |
| **CAST_N_T3_P2V2** | 3,008,736 | 2,994,015 | 2,979,353 | 2,388,688 | CCF | mWGS | | 2,467,263 |
| **CAST_N_T3_P2V4** | 3,635,261 | 3,625,577 | 3,402,188 | 3,150,852 | CCF | mWGS | | 2,467,263 |
| **CAST_R_T2_P1V2** | 2,060,652 | 2,044,533 | 1,790,501 | 1,481,878 | CCF | mWGS | | 1,347,724 |
| **CAST_R_T2_P1V4** | 2,568,781 | 2,559,562 | 2,263,056 | 1,867,295 | CCF | mWGS | | 1,347,724 |
| **CAST_R_T3_P2V2** | 3,074,781 | 3,060,900 | 2,875,804 | 2,400,294 | CCF | mWGS | | 2,072,473 |
| **CAST_R_T3_P2V4** | 4,651,224 | 4,640,804 | 4,363,023 | 4,062,138 | CCF | mWGS | | 2,072,473 |
| **CAST_R_T5_P2V2** | 1,552,076 | 1,542,807 | 1,534,954 | 1,245,932 | CCF | mWGS | | 1,257,428 |
| **CAST_R_T5_P2V4** | 1,947,227 | 1,936,456 | 1,757,833 | 1,589,676 | CCF | mWGS | | 1,257,428 |
| **PWK_2B_T1_P2V2** | 6,850,903 | 6,828,059 | 6,808,405 | 5,184,298 | CCF | mWGS | | 1,680,050 |
| **PWK_2B_T1_P2V4** | 4,700,651 | 4,690,531 | 4,235,591 | 3,694,563 | CCF | mWGS | | 1,680,050 |
| **PWK_2B_T3_P1V2** | 2,946,148 | 2,921,976 | 2,623,498 | 2,175,342 | CCF | mWGS | | 1,401,587 |
| **PWK_2B_T3_P1V4** | 1,541,417 | 1,533,867 | 1,387,641 | 1,185,499 | CCF | mWGS | | 1,401,587 |
| **PWK_2L_T1_P2V2** | 6,143,597 | 6,115,516 | 6,087,952 | 4,607,130 | CCF | mWGS | | 1,120,223 |
| **PWK_2L_T1_P2V4** | 5,876,098 | 5,858,957 | 5,338,608 | 4,650,757 | CCF | mWGS | | 1,120,223 |
| **PWK_2L_T3_P1V2** | 3,363,235 | 3,339,469 | 2,914,381 | 2,401,261 | CCF | mWGS | | 1,140,268 |

| | | | | | | | | |
|---|---|---|---|---|---|---|---|---|
| **PWK_2L_T3_P1V4** | 3,178,803 | 3,168,314 | 2,810,951 | 2,321,714 | CCF | mWGS | | 1,140,268 |
| **PWK_2L_T4_P2V2** | 6,687,291 | 6,655,413 | 6,617,380 | 4,895,269 | CCF | mWGS | | 2,231,733 |
| **PWK_2L_T4_P2V4** | 9,323,632 | 9,291,667 | 8,348,123 | 7,162,830 | CCF | mWGS | | 2,231,733 |
| **PWK_2R_T1_P2V2** | 3,761,962 | 3,752,641 | 3,741,317 | 2,766,857 | CCF | mWGS | | 1,409,132 |
| **PWK_2R_T1_P2V4** | 3,024,843 | 3,020,862 | 2,727,360 | 2,343,432 | CCF | mWGS | | 1,409,132 |
| **PWK_2R_T3_P2V2** | 3,857,441 | 3,846,854 | 3,374,060 | 2,730,550 | CCF | mWGS | | 1,219,049 |
| **PWK_2R_T3_P2V4** | 2,416,171 | 2,412,314 | 2,151,524 | 1,811,880 | CCF | mWGS | | 1,219,049 |
| **PWK_2R_T4_P2V2** | 3,486,917 | 3,476,494 | 3,462,740 | 2,508,432 | CCF | mWGS | | 1,954,048 |
| **PWK_2R_T4_P2V4** | 3,768,274 | 3,762,253 | 3,345,506 | 2,801,000 | CCF | mWGS | | 1,954,048 |
| **PWK_3R_T3_P2V2** | 2,079,710 | 2,050,161 | 1,856,245 | 1,554,683 | CCF | mWGS | | 1,191,034 |
| **PWK_3R_T3_P2V4** | 1,674,525 | 1,659,813 | 1,539,458 | 1,387,075 | CCF | mWGS | | 1,191,034 |
| **PWK_B_T1_P2V2** | 4,170,890 | 4,157,567 | 4,141,318 | 3,023,607 | CCF | mWGS | | 1,389,340 |
| **PWK_B_T1_P2V4** | 3,583,695 | 3,577,614 | 3,212,885 | 2,725,886 | CCF | mWGS | | 1,389,340 |
| **PWK_B_T4_P2V2** | 2,532,095 | 2,522,780 | 2,511,074 | 1,842,649 | CCF | mWGS | | 1,619,583 |
| **PWK_B_T4_P2V4** | 1,737,479 | 1,733,473 | 1,557,251 | 1,330,146 | CCF | mWGS | | 1,619,583 |
| **PWK_L_T3_P1V2** | 3,533,600 | 3,505,220 | 3,083,149 | 2,551,442 | CCF | mWGS | | 1,275,775 |
| **PWK_L_T3_P1V4** | 3,650,460 | 3,623,743 | 3,271,737 | 2,798,992 | CCF | mWGS | | 1,275,775 |
| **PWK_L_T4_P2V2** | 3,386,912 | 3,371,982 | 3,357,096 | 2,452,558 | CCF | mWGS | | 4,415,351 |
| **PWK_L_T4_P2V4** | 4,140,186 | 4,128,183 | 3,649,367 | 3,111,291 | CCF | mWGS | | 4,415,351 |
| **PWK_N_T1_P2V2** | 3,001,569 | 2,990,940 | 2,982,462 | 2,286,341 | CCF | mWGS | | 1,671,759 |
| **PWK_N_T1_P2V4** | 3,795,097 | 3,788,493 | 3,457,924 | 3,016,068 | CCF | mWGS | | 1,671,759 |
| **PWK_N_T3_P2V2** | 2,756,754 | 2,745,369 | 2,467,976 | 2,010,716 | CCF | mWGS | | 1,149,129 |
| **PWK_N_T3_P2V4** | 2,769,496 | 2,763,170 | 2,504,229 | 2,166,767 | CCF | mWGS | | 1,149,129 |
| **PWK_N_T4_P2V2** | 2,300,009 | 2,292,741 | 2,285,772 | 1,729,093 | CCF | mWGS | | 2,667,835 |
| **PWK_N_T4_P2V4** | 3,181,120 | 3,177,884 | 2,886,321 | 2,511,758 | CCF | mWGS | | 2,667,835 |
| **PWK_R_T1_P2V2** | 2,843,593 | 2,831,716 | 2,818,433 | 2,226,897 | CCF | mWGS | | 1,664,158 |
| **PWK_R_T1_P2V4** | 3,136,832 | 3,127,395 | 2,946,925 | 2,636,437 | CCF | mWGS | | 1,664,158 |
| **4M7_TD_P1V2** | 3,070,474 | 3,036,359 | 2,812,515 | 2,329,458 | HLB | mWGS, 16S | 13,843 | 14,536,240 |
| **4M7_TD_P1V4** | 4,217,339 | 4,158,973 | 3,777,222 | 3,369,637 | HLB | mWGS, 16S | 13,843 | 14,536,240 |
| **BF2_TD_P1V2** | 3,786,113 | 3,753,536 | 3,232,204 | 2,647,525 | HLB | mWGS, 16S | 17,306 | 17,562,416 |
| **BF2_TD_P1V4** | 3,703,215 | 3,679,057 | 3,203,312 | 2,877,310 | HLB | mWGS, 16S | 17,306 | 17,562,416 |
| **BF4_3D_P1V2** | 4,926,938 | 4,867,156 | 4,231,938 | 3,456,253 | HLB | mWGS, 16S | 23,858 | 20,657,550 |
| **BF4_3D_P1V4** | 7,419,491 | 7,325,177 | 6,303,220 | 5,578,788 | HLB | mWGS, 16S | 23,858 | 20,657,550 |
| **BM3_3D_P1V2** | 3,763,740 | 3,704,633 | 3,253,850 | 2,687,844 | HLB | mWGS, 16S | 18,739 | 16,036,400 |
| **BM3_3D_P1V4** | 2,540,999 | 2,485,541 | 2,162,079 | 1,921,752 | HLB | mWGS, 16S | 18,739 | 16,036,400 |
| **BF1_3D_P1V4** | 11,817,206 | 11,609,732 | 9,594,628 | 8,493,872 | HLB | mWGS, 16S | 19,112 | 16,128,555 |
| **4F6_3D_P1V2** | 2,279,215 | 2,262,079 | 2,087,399 | 1,741,066 | HLB | mWGS, 16S | 20,726 | 11,977,157 |
| **4F6_3D_P1V4** | 2,923,298 | 2,900,381 | 2,605,545 | 2,256,145 | HLB | mWGS, 16S | 20,726 | 11,977,157 |
| **4F7_0D_P1V2** | 4,108,993 | 4,050,220 | 3,887,878 | 3,260,226 | HLB | mWGS, 16S | 20,538 | 14,813,704 |
| **4F7_0D_P1V4** | 4,842,696 | 4,739,134 | 4,414,955 | 3,881,842 | HLB | mWGS, 16S | 20,538 | 14,813,704 |
| **4F7_TD_P1V2** | 2,749,784 | 2,717,855 | 2,499,177 | 2,065,976 | HLB | mWGS, 16S | 17,344 | 21,087,960 |
| **4F7_TD_P1V4** | 3,135,735 | 3,095,312 | 2,765,724 | 2,441,845 | HLB | mWGS, 16S | 17,344 | 21,087,960 |
| **4F8_0D_P1V2** | 3,680,925 | 3,647,783 | 3,565,836 | 2,992,261 | HLB | mWGS, 16S | 25,643 | 26,843,938 |
| **4F8_0D_P1V4** | 4,095,909 | 4,026,164 | 3,832,913 | 3,375,260 | HLB | mWGS, 16S | 25,643 | 26,843,938 |
| **4M1_0D_P1V2** | 2,988,400 | 2,912,627 | 2,777,891 | 2,335,125 | HLB | mWGS, 16S | 19,016 | 20,210,235 |
| **4M1_0D_P1V4** | 3,362,849 | 3,249,859 | 3,047,950 | 2,684,084 | HLB | mWGS, 16S | 19,016 | 20,210,235 |
| **4M1_TD_P1V2** | 1,886,222 | 1,842,969 | 1,694,802 | 1,386,882 | HLB | mWGS, 16S | 19,134 | 15,466,619 |
| **4M1_TD_P1V4** | 2,329,992 | 2,269,364 | 2,062,076 | 1,785,659 | HLB | mWGS, 16S | 19,134 | 15,466,619 |
| **4M3_0D_P1V2** | 2,243,817 | 2,179,653 | 2,029,830 | 1,682,699 | HLB | mWGS, 16S | 20,114 | 20,631,558 |
| **4M3_0D_P1V4** | 3,391,841 | 3,249,050 | 2,953,131 | 2,542,524 | HLB | mWGS, 16S | 20,114 | 20,631,558 |
| **4M3_TD_P1V2** | 1,941,052 | 1,905,829 | 1,780,845 | 1,500,845 | HLB | mWGS, 16S | N/A | 12,800,828 |
| **4M3_TD_P1V4** | 2,734,471 | 2,670,658 | 2,474,377 | 2,204,178 | HLB | mWGS, 16S | N/A | 12,800,828 |
| **4M5_0D_P1V2** | 2,885,483 | 2,845,952 | 2,737,956 | 2,258,034 | HLB | mWGS, 16S | 17,908 | 9,206,898 |
| **4M5_0D_P1V4** | 6,109,395 | 5,955,736 | 5,510,834 | 4,924,147 | HLB | mWGS, 16S | 17,908 | 9,206,898 |
| **4M6_0D_P1V2** | 3,282,804 | 3,260,945 | 3,113,238 | 2,608,554 | HLB | mWGS, 16S | 16,261 | 11,791,275 |
| **4M6_0D_P1V4** | 4,735,775 | 4,675,061 | 4,296,485 | 3,875,883 | HLB | mWGS, 16S | 16,261 | 11,791,275 |

| | | | | | | | | |
|---|---|---|---|---|---|---|---|---|
| **4M6_TD_P1V2** | 2,056,491 | 2,039,847 | 1,896,313 | 1,568,065 | HLB | mWGS, 16S | 14,840 | 12,270,221 |
| **4M6_TD_P1V4** | 4,001,805 | 3,963,407 | 3,640,111 | 3,217,749 | HLB | mWGS, 16S | 14,840 | 12,270,221 |
| **4M7_0D_P1V2** | 2,899,401 | 2,873,617 | 2,741,280 | 2,282,043 | HLB | mWGS, 16S | 14,413 | 12,370,629 |
| **4M7_0D_P1V4** | 5,252,191 | 5,148,391 | 4,683,041 | 4,196,176 | HLB | mWGS, 16S | 14,413 | 12,370,629 |
| **4M7_3D_P1V2** | 332,738 | 327,480 | 298,705 | 250,132 | HLB | mWGS, 16S | 21,708 | 12,816,146 |
| **4M7_3D_P1V4** | 3,104,676 | 3,052,663 | 2,754,386 | 2,432,690 | HLB | mWGS, 16S | 21,708 | 12,816,146 |
| **BF1_3D_P1V2** | 4,349,421 | 4,290,670 | 3,608,813 | 2,918,302 | HLB | mWGS, 16S | 19,112 | 16,128,555 |
| **BF1_TD_P1V2** | 4,349,084 | 4,301,643 | 3,718,700 | 3,077,284 | HLB | mWGS, 16S | 15,258 | 19,063,564 |
| **BF1_TD_P1V4** | 10,818,241 | 10,725,494 | 9,395,150 | 8,412,329 | HLB | mWGS, 16S | 15,258 | 19,063,564 |
| **BF2_0D_P1V2** | 1,036,862 | 1,022,884 | 990,237 | 822,017 | HLB | mWGS, 16S | 16,350 | 16,094,974 |
| **BF2_0D_P1V4** | 2,887,509 | 2,833,867 | 2,716,200 | 2,459,640 | HLB | mWGS, 16S | 16,350 | 16,094,974 |
| **BF2_3D_P1V2** | 4,463,930 | 4,408,625 | 3,922,982 | 3,187,687 | HLB | mWGS, 16S | 20,352 | 16,996,156 |
| **BF2_3D_P1V4** | 5,144,660 | 5,080,285 | 4,468,054 | 3,948,631 | HLB | mWGS, 16S | 20,352 | 16,996,156 |
| **BF3_0D_P1V2** | 1,076,265 | 1,059,622 | 1,009,364 | 839,454 | HLB | mWGS, 16S | 14,604 | 4,173,812 |
| **BF3_0D_P1V4** | 1,480,319 | 1,452,153 | 1,378,522 | 1,250,574 | HLB | mWGS, 16S | 14,604 | 4,173,812 |
| **BF3_TD_P1V2** | 3,886,155 | 3,835,072 | 3,230,264 | 2,617,684 | HLB | mWGS, 16S | 19,094 | 22,040,550 |
| **BF3_TD_P1V4** | 6,609,591 | 6,554,349 | 5,651,804 | 5,047,968 | HLB | mWGS, 16S | 19,094 | 22,040,550 |
| **BM1_0D_P1V2** | 405,147 | 398,951 | 384,314 | 319,025 | HLB | mWGS, 16S | 20,775 | 2,145,125 |
| **BM1_0D_P1V4** | 908,874 | 894,866 | 857,860 | 777,637 | HLB | mWGS, 16S | 20,775 | 2,145,125 |
| **BM1_3D_P1V2** | 3,959,852 | 3,932,255 | 3,631,267 | 3,087,740 | HLB | mWGS, 16S | 21,829 | 17,036,259 |
| **BM1_3D_P1V4** | 3,554,852 | 3,524,837 | 3,263,947 | 2,924,582 | HLB | mWGS, 16S | 21,829 | 17,036,259 |
| **BM1_TD_P1V2** | 3,780,907 | 3,724,486 | 3,290,011 | 2,705,698 | HLB | mWGS, 16S | 20,138 | 16,170,189 |
| **BM1_TD_P1V4** | 5,595,070 | 5,493,890 | 4,848,802 | 4,314,952 | HLB | mWGS, 16S | 20,138 | 16,170,189 |
| **BM2_0D_P1V2** | 470,614 | 461,780 | 442,711 | 368,151 | HLB | mWGS, 16S | 18,099 | 380,675 |
| **BM2_0D_P1V4** | 601,847 | 581,695 | 544,661 | 494,357 | HLB | mWGS, 16S | 18,099 | 380,675 |
| **BM2_TD_P1V2** | 6,646,497 | 6,566,074 | 5,777,723 | 4,772,685 | HLB | mWGS, 16S | 21,880 | 19,304,612 |
| **BM2_TD_P1V4** | 7,952,150 | 7,843,074 | 6,941,847 | 6,218,731 | HLB | mWGS, 16S | 21,880 | 19,304,612 |
| **BM3_0D_P1V2** | 8,031,520 | 7,912,289 | 7,497,222 | 6,158,660 | HLB | mWGS, 16S | 20,054 | 18,817,236 |
| **BM3_0D_P1V4** | 8,541,347 | 8,443,523 | 7,974,767 | 7,187,515 | HLB | mWGS, 16S | 20,054 | 18,817,236 |
| **A_2_F01A_P2V4** | 3,391,647 | 3,382,935 | 3,311,152 | 3,005,214 | VNDR | 16S | 15,441 | |
| **A_1_M46A_P2V2** | 2,161,998 | 2,134,107 | 1,945,192 | 1,635,696 | VNDR | 16S | 22,385 | |
| **A_1_M46A _P2V4** | 2,736,372 | 2,706,418 | 2,424,260 | 2,128,711 | VNDR | 16S | 22,385 | |
| **A_2_F01A _P2V2** | 1,989,779 | 1,983,378 | 1,924,039 | 1,609,975 | VNDR | 16S | 15,441 | |
| **B_3_M51B_P2V2** | 2,973,987 | 2,961,913 | 2,753,108 | 2,462,391 | VNDR | 16S | 19,170 | |
| **B_3_M51B _P2V4** | 4,285,064 | 4,274,413 | 4,038,946 | 3,797,165 | VNDR | 16S | 19,170 | |
| **Ecoli_P1V2** | 2,735,448 | 2,732,889 | 2,157,203 | 2,086,420 | This study | | | |
| **ecoli_P2V2** | 5,526,780 | 5,521,930 | 4,417,647 | 4,257,598 | This study | | | |
| **ecoli_P1V4** | 4,063,753 | 4,058,214 | 2,420,869 | 1,924,636 | This study | | | |
| **ecoli_P2V4** | 6,183,444 | 6,178,204 | 3,833,500 | 3,103,533 | This study | | | |
| **human_stool_P1V2** | 328,929 | 324,202 | 289,074 | 260,278 | This study | | | |
| **human_stool_P2V2** | 277,616 | 273,845 | 243,484 | 218,040 | This study | | | |
| **human_stool_P1V4** | 452,413 | 450,956 | 380,204 | 352,381 | This study | | | |
| **human_stool_P2V4** | 576,879 | 575,295 | 484,286 | 452,870 | This study | | | |
| **zymo_mock_P1V2** | 1,506,453 | 1,502,222 | 1,277,278 | 1,179,509 | This study | | | |
| **zymo_mock_P2V2** | 1,324,645 | 1,321,668 | 1,131,085 | 1,039,500 | This study | | | |
| **zymo_mock_P1V4** | 848,994 | 846,842 | 564,505 | 464,554 | This study | | | |
| **zymo_mock_P2V4** | 1,809,909 | 1,806,699 | 1,228,518 | 1,019,616 | This study | | | |
| **NTC_CCF_VNDR_KOMP_P2V2** | 747 | 745 | 632 | 473 | This study | | | |
| **NTC_CCF_VNDR_KOMP_P2V4** | 3,649 | 3,648 | 3,648 | 3,545 | This study | | | |
| **NTC_HL44_P1V2** | 592 | 572 | 533 | 468 | This study | | | |
| **NTC_HL44_P1V4** | 3,416 | 3,403 | 3,391 | 3,178 | This study | | | |
