## Supplemental Table 4 for "Genome-based targeted sequencing as a reproducible microbial community profiling assay"

| | | | HLB vs. B6 (HF) | | | | | | HLB vs. B6 (Chow) | | | | | | HLB (Chow vs. HF) | | | | | | B6 (Chow vs. HF) | | | | | |
|---|---|---|---|---|---|---|---|---|---|---|---|---|---|---|---|---|---|---|---|---|---|---|---|---|---|---|
| | | | **Allegro** | | | **JAX** | | | **Allegro** | | | **JAX** | | | **Allegro** | | | **JAX** | | | **Allegro** | | | **JAX** | | |
| **KO** | **Pathway Class** | **Pathway** | **Class** | **LDA** | **p-value** | **Class** | **LDA** | **p-value** | **Class** | **LDA** | **p-value** | **Class** | **LDA** | **p-value** | **Class** | **LDA** | **p-value** | **Class** | **LDA** | **p-value** | **Class** | **LDA** | **p-value** | **Class** | **LDA** | **p-value** |
| **ko00400** | Amino Acid Metabolism | Phenylalanine, tyrosine and tryptophan biosynthesis | B6 | -2.045 | 0.023 | B6 | -2.395 | 0.005 | - | - | - | - | - | - | Chow | -2.18 | 0.015 | Chow | -2.203 | 0.022 | - | - | - | - | - | - |
| **ko00290** | Amino Acid Metabolism | Valine, leucine and isoleucine biosynthesis | B6 | -2.022 | 0.023 | B6 | -2.072 | 0.03 | B6 | -2.515 | 0.004 | B6 | -2.476 | 0.004 | HF | 2.617 | 0.003 | HF | 2.597 | 0.003 | HF | 2.252 | 0.002 | HF | 2.299 | 0.002 |
| **ko00220** | Amino Acid Metabolism | Arginine biosynthesis | B6 | -2.158 | 0.007 | B6 | -2.13 | 0.002 | - | - | - | - | - | - | Chow | -2.363 | 0.007 | Chow | -2.308 | 0.01 | - | - | - | - | - | - |
| **ko00471** | Amino Acid Metabolism | D-Glutamine and D-glutamate metabolism | B6 | -2.097 | 0.001 | B6 | -2.203 | 0.001 | HLB444 | 2.317 | 0.004 | HLB444 | 2.287 | 0.004 | Chow | -2.488 | 0.003 | Chow | -2.494 | 0.003 | - | - | - | - | - | - |
| **ko00450** | Amino Acid Metabolism | Selenocompound metabolism | HLB444 | 2.241 | 0.001 | HLB444 | 2.303 | 0.001 | - | - | - | B6 | -2.015 | 0.004 | HF | 2.455 | 0.003 | HF | 2.456 | 0.003 | - | - | - | - | - | - |
| **ko00270** | Amino Acid Metabolism | Cysteine and methionine metabolism | HLB444 | 2.345 | 0.001 | HLB444 | 2.345 | 0.001 | - | - | - | - | - | - | HF | 2.552 | 0.003 | HF | 2.54 | 0.003 | HF | 2.243 | 0.002 | HF | 2.241 | 0.01 |
| **ko00300** | Amino Acid Metabolism | Lysine biosynthesis | - | - | - | - | - | - | B6 | -2.192 | 0.004 | B6 | -2.218 | 0.019 | HF | 2.466 | 0.003 | HF | 2.434 | 0.003 | HF | 2.341 | 0.002 | HF | 2.331 | 0.002 |
| **ko00350** | Amino Acid Metabolism | Tyrosine metabolism | - | - | - | - | - | - | B6 | -2.217 | 0.004 | B6 | -2.167 | 0.004 | HF | 2.439 | 0.003 | HF | 2.42 | 0.003 | HF | 2.195 | 0.002 | HF | 2.247 | 0.002 |
| **ko00340** | Amino Acid Metabolism | Histidine metabolism | - | - | - | - | - | - | B6 | -2.209 | 0.012 | - | - | - | HF | 2.482 | 0.003 | HF | 2.468 | 0.003 | HF | 2.23 | 0.002 | HF | 2.292 | 0.002 |
| **ko00250** | Amino Acid Metabolism | Alanine, aspartate and glutamate metabolism | - | - | - | - | - | - | HLB444 | 2.532 | 0.004 | HLB444 | 2.516 | 0.004 | Chow | -2.82 | 0.003 | Chow | -2.797 | 0.003 | Chow | -2.483 | 0.002 | Chow | -2.586 | 0.002 |
| **ko00260** | Amino Acid Metabolism | Glycine, serine and threonine metabolism | - | - | - | B6 | -2.124 | 0.002 | HLB444 | 2.084 | 0.028 | HLB444 | 2.318 | 0.012 | - | - | - | - | - | - | HF | 2.351 | 0.002 | HF | 2.524 | 0.002 |
| **ko00430** | Amino Acid Metabolism | Taurine and hypotaurine metabolism | - | - | - | - | - | - | HLB444 | 2.256 | 0.004 | HLB444 | 2.251 | 0.007 | Chow | -2.409 | 0.003 | Chow | -2.39 | 0.003 | Chow | -2.108 | 0.002 | Chow | -2.171 | 0.002 |
| **ko00280** | Amino Acid Metabolism | Valine, leucine and isoleucine degradation | - | - | - | - | - | - | HLB444 | 2.546 | 0.012 | HLB444 | 2.643 | 0.007 | Chow | -2.688 | 0.004 | Chow | -2.659 | 0.004 | - | - | - | - | - | - |
| **ko00410** | Amino Acid Metabolism | beta-Alanine metabolism | - | - | - | - | - | - | HLB444 | 2.055 | 0.004 | HLB444 | 2.147 | 0.004 | Chow | -2.261 | 0.003 | Chow | -2.248 | 0.003 | - | - | - | - | - | - |
| **ko00330** | Amino Acid Metabolism | Arginine and proline metabolism | - | - | - | - | - | - | HLB444 | 2.254 | 0.007 | HLB444 | 2.372 | 0.004 | Chow | -2.562 | 0.003 | Chow | -2.554 | 0.003 | Chow | -2.282 | 0.002 | - | - | - |
| **ko00460** | Amino Acid Metabolism | Cyanoamino acid metabolism | - | - | - | HLB444 | 2.07 | 0.005 | - | - | - | - | - | - | Chow | -2.212 | 0.01 | Chow | -2.196 | 0.01 | Chow | -2.484 | 0.002 | Chow | -2.536 | 0.002 |
| **ko00440** | Amino Acid Metabolism | Phosphonate and phosphinate metabolism | - | - | - | - | - | - | - | - | - | - | - | - | HF | 2.213 | 0.003 | HF | 2.184 | 0.003 | HF | 2.075 | 0.002 | HF | 2.036 | 0.002 |
| **ko00310** | Amino Acid Metabolism | Lysine degradation | - | - | - | - | - | - | - | - | - | - | - | - | HF | 2.315 | 0.003 | HF | 2.305 | 0.003 | HF | 2.183 | 0.002 | HF | 2.255 | 0.002 |
| **ko00360** | Amino Acid Metabolism | Phenylalanine metabolism | - | - | - | - | - | - | - | - | - | - | - | - | HF | 2.205 | 0.003 | HF | 2.212 | 0.003 | - | - | - | - | - | - |
| **ko00480** | Amino Acid Metabolism | Glutathione metabolism | - | - | - | HLB444 | 2.034 | 0.002 | - | - | - | - | - | - | HF | 2.118 | 0.004 | HF | 2.13 | 0.004 | - | - | - | - | - | - |
| **ko00380** | Amino Acid Metabolism | Tryptophan metabolism | - | - | - | - | - | - | - | - | - | - | - | - | - | - | - | - | - | - | - | - | - | HF | 2.076 | 0.002 |
| **ko01130** | Biosynthesis of antibiotics | Biosynthesis of antibiotics | B6 | -2.591 | 0.002 | B6 | -2.708 | 0.001 | - | - | - | - | - | - | - | - | - | - | - | - | HF | 2.358 | 0.037 | HF | 2.656 | 0.002 |
| **ko00521** | Biosynthesis of other Secondary Metabolites | Streptomycin biosynthesis | B6 | -2.07 | 0.017 | - | - | - | - | - | - | HLB444 | 2.057 | 0.028 | Chow | -2.526 | 0.003 | Chow | -2.493 | 0.003 | Chow | -2.191 | 0.002 | Chow | -2.133 | 0.005 |
| **ko00401** | Biosynthesis of other Secondary Metabolites | Novobiocin biosynthesis | - | - | - | - | - | - | B6 | -2.035 | 0.004 | - | - | - | HF | 2.339 | 0.003 | HF | 2.323 | 0.003 | HF | 2.208 | 0.002 | HF | 2.289 | 0.002 |

| | | | | | | | | | | | | | | | | | | | | | | | | | | |
|---|---|---|---|---|---|---|---|---|---|---|---|---|---|---|---|---|---|---|---|---|---|---|---|---|---|---|
| **ko00525** | Biosynthesis of other Secondary Metabolites | Acarbose and validamycin biosynthesis | - | - | - | - | - | - | HLB444 | 2.011 | 0.028 | HLB444 | 2.1 | 0.019 | Chow | -2.568 | 0.003 | Chow | -2.554 | 0.003 | Chow | -2.3 | 0.002 | Chow | -2.28 | 0.002 |
| **ko00901** | Biosynthesis of other Secondary Metabolites | Indole alkaloid biosynthesis | - | - | - | - | - | - | HLB444 | 2.117 | 0.015 | HLB444 | 2.195 | 0.004 | - | - | - | - | - | - | - | - | - | - | - | - |
| **ko00311** | Biosynthesis of other Secondary Metabolites | Penicillin and cephalosporin biosynthesis | - | - | - | - | - | - | - | - | - | - | - | - | Chow | -2.053 | 0.003 | Chow | -2.006 | 0.003 | - | - | - | - | - | - |
| **ko00940** | Biosynthesis of other Secondary Metabolites | Phenylpropanoid biosynthesis | - | - | - | HLB444 | 2.132 | 0.001 | - | - | - | B6 | -2.029 | 0.019 | Chow | -2.228 | 0.003 | Chow | -2.221 | 0.003 | Chow | -2.56 | 0.002 | Chow | -2.61 | 0.002 |
| **ko00524** | Biosynthesis of other Secondary Metabolites | Neomycin, kanamycin and gentamicin biosynthesis | - | - | - | - | - | - | - | - | - | - | - | - | HF | 2.136 | 0.003 | HF | 2.133 | 0.003 | HF | 2.132 | 0.002 | HF | 2.091 | 0.002 |
| **ko00960** | Biosynthesis of other Secondary Metabolites | Tropane, piperidine and pyridine alkaloid biosynthesis | - | - | - | - | - | - | - | - | - | - | - | - | HF | 2.133 | 0.003 | HF | 2.111 | 0.003 | HF | 2.005 | 0.002 | HF | 2.08 | 0.002 |
| **ko00232** | Biosynthesis of other Secondary Metabolites | Caffeine metabolism | - | - | - | - | - | - | - | - | - | - | - | - | HF | 2.181 | 0.003 | HF | 2.238 | 0.003 | - | - | - | - | - | - |
| **ko00261** | Biosynthesis of other Secondary Metabolites | Monobactam biosynthesis | - | - | - | - | - | - | - | - | - | B6 | -2.063 | 0.004 | - | - | - | - | - | - | - | - | - | - | - | - |
| **ko00620** | Carbohydrate Metabolism | Pyruvate metabolism | B6 | -2.132 | 0.005 | B6 | -2.174 | 0.002 | HLB444 | 2.422 | 0.004 | HLB444 | 2.467 | 0.004 | Chow | -2.18 | 0.022 | Chow | -2.138 | 0.032 | HF | 2.385 | 0.002 | HF | 2.473 | 0.002 |
| **ko00020** | Carbohydrate Metabolism | Citrate cycle (TCA cycle) | B6 | -2.624 | 0.001 | B6 | -2.674 | 0.001 | HLB444 | 2.334 | 0.042 | HLB444 | 2.529 | 0.028 | Chow | -2.736 | 0.004 | Chow | -2.736 | 0.004 | - | - | - | HF | 2.343 | 0.027 |
| **ko00500** | Carbohydrate Metabolism | Starch and sucrose metabolism | HLB444 | 2.826 | 0.001 | HLB444 | 2.89 | 0.001 | HLB444 | 2.39 | 0.028 | - | - | - | Chow | -2.781 | 0.003 | Chow | -2.728 | 0.015 | Chow | -3.002 | 0.002 | Chow | -3.096 | 0.002 |
| **ko00040** | Carbohydrate Metabolism | Pentose and glucuronate interconversions | HLB444 | 2.478 | 0.001 | HLB444 | 2.622 | 0.001 | B6 | -2.84 | 0.004 | B6 | -2.863 | 0.004 | - | - | - | - | - | - | Chow | -3.049 | 0.002 | Chow | -3.105 | 0.002 |
| **ko00520** | Carbohydrate Metabolism | Amino sugar and nucleotide sugar metabolism | HLB444 | 2.673 | 0.003 | HLB444 | 2.752 | 0.002 | B6 | -2.649 | 0.004 | B6 | -2.686 | 0.007 | - | - | - | - | - | - | Chow | -2.869 | 0.002 | Chow | -2.932 | 0.002 |
| **ko00052** | Carbohydrate Metabolism | Galactose metabolism | HLB444 | 2.316 | 0.007 | HLB444 | 2.246 | 0.017 | HLB444 | 2.393 | 0.004 | HLB444 | 2.318 | 0.004 | Chow | -2.359 | 0.032 | Chow | -2.351 | 0.046 | Chow | -2.28 | 0.002 | Chow | -2.27 | 0.002 |
| **ko00051** | Carbohydrate Metabolism | Fructose and mannose metabolism | HLB444 | 2.666 | 0.002 | HLB444 | 2.634 | 0.001 | HLB444 | 2.407 | 0.004 | HLB444 | 2.315 | 0.004 | Chow | -2.803 | 0.003 | Chow | -2.748 | 0.003 | Chow | -2.929 | 0.002 | Chow | -2.894 | 0.002 |
| **ko00030** | Carbohydrate Metabolism | Pentose phosphate pathway | HLB444 | 2.634 | 0.001 | HLB444 | 2.705 | 0.001 | B6 | -2.408 | 0.019 | B6 | -2.548 | 0.012 | HF | 2.879 | 0.003 | HF | 2.857 | 0.003 | - | - | - | - | - | - |
| **ko00562** | Carbohydrate Metabolism | Inositol phosphate metabolism | - | - | - | - | - | - | B6 | -2.08 | 0.004 | B6 | -2.012 | 0.004 | HF | 2.108 | 0.015 | HF | 2.181 | 0.003 | - | - | - | - | - | - |
| **ko00660** | Carbohydrate Metabolism | C5-Branched dibasic acid metabolism | - | - | - | - | - | - | B6 | -2.352 | 0.007 | B6 | -2.344 | 0.012 | HF | 2.51 | 0.003 | HF | 2.497 | 0.003 | HF | 2.065 | 0.003 | HF | 2.064 | 0.007 |
| **ko00010** | Carbohydrate Metabolism | Glycolysis / Gluconeogenesis | - | - | - | - | - | - | HLB444 | 2.152 | 0.004 | HLB444 | 2.054 | 0.004 | HF | 2.504 | 0.003 | HF | 2.543 | 0.003 | HF | 2.57 | 0.002 | HF | 2.572 | 0.002 |
| **ko00640** | Carbohydrate Metabolism | Propanoate metabolism | - | - | - | - | - | - | HLB444 | 2.078 | 0.028 | - | - | - | - | - | - | - | - | - | - | - | - | - | - | - |
| **ko00630** | Carbohydrate Metabolism | Glyoxylate and dicarboxylate metabolism | - | - | - | B6 | -2.086 | 0.039 | HLB444 | 2.228 | 0.028 | HLB444 | 2.414 | 0.019 | Chow | -2.729 | 0.003 | Chow | -2.726 | 0.003 | Chow | -2.471 | 0.002 | Chow | -2.252 | 0.007 |
| **ko00053** | Carbohydrate Metabolism | Ascorbate and aldarate metabolism | - | - | - | - | - | - | - | - | - | - | - | - | - | - | - | - | - | - | Chow | -2.209 | 0.002 | Chow | -2.18 | 0.002 |
| **ko00650** | Carbohydrate Metabolism | Butanoate metabolism | - | - | - | - | - | - | - | - | - | - | - | - | - | - | - | - | - | - | - | - | - | Chow | -2.161 | 0.003 |
| **ko02040** | Cell Motility | Flagellar assembly | HLB444 | 3.062 | 0.002 | HLB444 | 3.132 | 0.002 | B6 | -2.77 | 0.028 | B6 | -2.879 | 0.028 | HF | 3.292 | 0.003 | HF | 3.288 | 0.004 | - | - | - | - | - | - |
| **ko02030** | Cell Motility | Bacterial chemotaxis | HLB444 | 3.182 | 0.001 | HLB444 | 3.245 | 0.001 | B6 | -2.822 | 0.028 | B6 | -2.966 | 0.028 | HF | 3.284 | 0.004 | HF | 3.286 | 0.004 | - | - | - | Chow | -2.809 | 0.014 |
| **ko02025** | Cellular Community | Biofilm formation - Pseudomonas aeruginosa | HLB444 | 2.099 | 0.002 | HLB444 | 2.215 | 0.001 | - | - | - | - | - | - | - | - | - | - | - | - | Chow | -2.185 | 0.002 | Chow | -2.281 | 0.002 |

| | | | | | | | | | | | | | | | | | | | | | | | | | | |
|---|---|---|---|---|---|---|---|---|---|---|---|---|---|---|---|---|---|---|---|---|---|---|---|---|---|---|
| **ko02024** | Cellular Community | Quorum sensing | HLB444 | 2.858 | 0.001 | HLB444 | 2.83 | 0.003 | B6 | -2.578 | 0.028 | B6 | -2.783 | 0.012 | HF | 3.11 | 0.003 | HF | 3.101 | 0.003 | - | - | - | - | - | - |
| **ko02026** | Cellular Community | Biofilm formation - Escherichia coli | HLB444 | 2.286 | 0.001 | HLB444 | 2.393 | 0.001 | - | - | - | - | - | - | - | - | - | - | - | - | Chow | -2.422 | 0.002 | Chow | -2.539 | 0.002 |
| **ko01503** | Drug Resistance: Antimicrobial | Cationic antimicrobial peptide (CAMP) resistance | B6 | -2.588 | 0.001 | B6 | -2.703 | 0.001 | - | - | - | - | - | - | Chow | -2.649 | 0.003 | Chow | -2.651 | 0.004 | - | - | - | - | - | - |
| **ko01501** | Drug Resistance: Antimicrobial | beta-Lactam resistance | B6 | -2.492 | 0.002 | B6 | -2.633 | 0.001 | B6 | -2.492 | 0.004 | B6 | -2.484 | 0.004 | HF | 2.497 | 0.003 | HF | 2.501 | 0.003 | HF | 2.472 | 0.002 | HF | 2.623 | 0.002 |
| **ko01502** | Drug Resistance: Antimicrobial | Vancomycin resistance | - | - | - | B6 | -2.035 | 0.003 | - | - | - | - | - | - | HF | 2.325 | 0.004 | HF | 2.28 | 0.004 | HF | 2.414 | 0.002 | HF | 2.408 | 0.002 |
| **ko01523** | Drug Resistance: Antineoplastic | Antifolate resistance | - | - | - | - | - | - | HLB444 | 2.154 | 0.004 | HLB444 | 2.131 | 0.004 | - | - | - | - | - | - | - | - | - | - | - | - |
| **ko03320** | Endocrine System | PPAR signaling pathway | - | - | - | - | - | - | - | - | - | - | - | - | Chow | -2.442 | 0.003 | Chow | -2.437 | 0.003 | Chow | -2.234 | 0.002 | Chow | -2.242 | 0.002 |
| **ko00190** | Energy Metabolism | Oxidative phosphorylation | B6 | -2.862 | 0.001 | B6 | -2.967 | 0.001 | HLB444 | 2.703 | 0.028 | HLB444 | 2.8 | 0.028 | Chow | -3.092 | 0.004 | Chow | -3.1 | 0.004 | - | - | - | - | - | - |
| **ko00720** | Energy Metabolism | Carbon fixation pathways in prokaryotes | B6 | -2.169 | 0.039 | - | - | - | HLB444 | 2.576 | 0.019 | HLB444 | 2.66 | 0.012 | Chow | -2.72 | 0.007 | Chow | -2.68 | 0.007 | - | - | - | - | - | - |
| **ko00920** | Energy Metabolism | Sulfur metabolism | HLB444 | 2.364 | 0.001 | HLB444 | 2.419 | 0.001 | B6 | -2.089 | 0.028 | B6 | -2.25 | 0.019 | HF | 2.614 | 0.003 | HF | 2.608 | 0.003 | - | - | - | - | - | - |
| **ko00710** | Energy Metabolism | Carbon fixation in photosynthetic organisms | HLB444 | 2.054 | 0.009 | HLB444 | 2.135 | 0.002 | - | - | - | - | - | - | Chow | -2.047 | 0.032 | Chow | -2.025 | 0.032 | Chow | -2.106 | 0.002 | Chow | -2.187 | 0.002 |
| **ko00680** | Energy Metabolism | Methane metabolism | HLB444 | 2.468 | 0.001 | HLB444 | 2.531 | 0.001 | HLB444 | 2.113 | 0.019 | HLB444 | 2.146 | 0.019 | - | - | - | HF | 2.182 | 0.032 | - | - | - | - | - | - |
| **ko00195** | Energy Metabolism | Photosynthesis | - | - | - | - | - | - | HLB444 | 2.081 | 0.012 | - | - | - | Chow | -2.394 | 0.003 | Chow | -2.396 | 0.003 | - | - | - | Chow | -2.05 | 0.002 |
| **ko00910** | Energy Metabolism | Nitrogen metabolism | - | - | - | HLB444 | 2.024 | 0.013 | - | - | - | - | - | - | Chow | -2.168 | 0.01 | Chow | -2.127 | 0.007 | Chow | -2.285 | 0.002 | Chow | -2.438 | 0.002 |
| **ko03060** | Folding, Sorting, and Degradation | Protein export | - | - | - | - | - | - | HLB444 | 2.025 | 0.004 | - | - | - | Chow | -2.267 | 0.003 | Chow | -2.265 | 0.003 | Chow | -2.185 | 0.002 | Chow | -2.208 | 0.002 |
| **ko03018** | Folding, Sorting, and Degradation | RNA degradation | - | - | - | B6 | -2.044 | 0.03 | HLB444 | 2.179 | 0.007 | HLB444 | 2.111 | 0.028 | Chow | -2.081 | 0.022 | - | - | - | - | - | - | HF | 2.037 | 0.01 |
| **ko01100** | Global Overview Maps | Metabolic pathways | B6 | -3.314 | 0.003 | B6 | -3.324 | 0.003 | HLB444 | 3.221 | 0.028 | HLB444 | 3.343 | 0.028 | Chow | -3.789 | 0.003 | Chow | -3.784 | 0.003 | Chow | -3.41 | 0.002 | Chow | -3.289 | 0.002 |
| **ko01110** | Global Overview Maps | Biosynthesis of secondary metabolites | B6 | -2.904 | 0.001 | B6 | -3.012 | 0.001 | - | - | - | - | - | - | - | - | - | - | - | - | - | - | - | HF | 2.825 | 0.002 |
| **ko01120** | Global Overview Maps | Microbial metabolism in diverse environments | - | - | - | - | - | - | B6 | -2.592 | 0.019 | B6 | -2.501 | 0.019 | HF | 3.272 | 0.003 | HF | 3.247 | 0.003 | HF | 3.192 | 0.002 | HF | 3.158 | 0.002 |
| **ko01230** | Global Overview Maps | Biosynthesis of amino acids | - | - | - | - | - | - | B6 | -2.976 | 0.007 | B6 | -2.979 | 0.007 | HF | 3.326 | 0.003 | HF | 3.305 | 0.003 | HF | 3.004 | 0.002 | HF | 3.005 | 0.002 |
| **ko01210** | Global Overview Maps | 2-Oxocarboxylic acid metabolism | - | - | - | - | - | - | B6 | -2.631 | 0.004 | B6 | -2.574 | 0.007 | HF | 2.919 | 0.003 | HF | 2.905 | 0.003 | HF | 2.589 | 0.002 | HF | 2.653 | 0.002 |
| **ko01220** | Global Overview Maps | Degradation of aromatic compounds | - | - | - | - | - | - | B6 | -2.446 | 0.004 | B6 | -2.416 | 0.004 | HF | 2.445 | 0.003 | HF | 2.419 | 0.003 | - | - | - | - | - | - |
| **ko01200** | Global Overview Maps | Carbon metabolism | - | - | - | - | - | - | HLB444 | 2.648 | 0.004 | HLB444 | 2.701 | 0.004 | HF | 2.525 | 0.046 | HF | 2.508 | 0.046 | HF | 2.804 | 0.002 | HF | 2.792 | 0.002 |
| **ko01212** | Global Overview Maps | Fatty acid metabolism | - | - | - | - | - | - | - | - | - | - | - | - | Chow | -2.431 | 0.003 | Chow | -2.407 | 0.003 | Chow | -2.228 | 0.014 | Chow | -2.243 | 0.01 |
| **ko00604** | Glycan Biosynthesis and Metabolism | Glycosphingolipid biosynthesis - ganglio series | B6 | -2.542 | 0.001 | B6 | -2.598 | 0.001 | B6 | -2.084 | 0.004 | - | - | - | HF | 2.144 | 0.032 | HF | 2.06 | 0.032 | HF | 2.533 | 0.002 | HF | 2.612 | 0.002 |
| **ko00513** | Glycan Biosynthesis and Metabolism | Various types of N-glycan biosynthesis | B6 | -2.55 | 0.001 | B6 | -2.604 | 0.001 | B6 | -2.094 | 0.004 | - | - | - | HF | 2.143 | 0.032 | HF | 2.058 | 0.032 | HF | 2.537 | 0.002 | HF | 2.614 | 0.002 |
| **ko00511** | Glycan Biosynthesis and Metabolism | Other glycan degradation | B6 | -2.916 | 0.001 | B6 | -2.957 | 0.001 | B6 | -2.501 | 0.007 | - | - | - | - | - | - | - | - | - | HF | 2.739 | 0.002 | HF | 2.855 | 0.002 |

| | | | | | | | | | | | | | | | | | | | | | | | | | | |
|---|---|---|---|---|---|---|---|---|---|---|---|---|---|---|---|---|---|---|---|---|---|---|---|---|---|---|
| **ko00531** | Glycan Biosynthesis and Metabolism | Glycosaminoglycan degradation | B6 | -2.685 | 0.001 | B6 | -2.75 | 0.001 | B6 | -2.319 | 0.004 | B6 | -2.117 | 0.007 | HF | 2.499 | 0.003 | HF | 2.437 | 0.01 | HF | 2.741 | 0.002 | HF | 2.819 | 0.002 |
| **ko00603** | Glycan Biosynthesis and Metabolism | Glycosphingolipid biosynthesis - globo and isoglobo series | B6 | -2.548 | 0.001 | B6 | -2.634 | 0.001 | B6 | -2.012 | 0.004 | - | - | - | HF | 2.147 | 0.022 | HF | 2.039 | 0.032 | HF | 2.565 | 0.002 | HF | 2.645 | 0.002 |
| **ko00540** | Glycan Biosynthesis and Metabolism | Lipopolysaccharide biosynthesis | B6 | -2.746 | 0.002 | B6 | -2.826 | 0.002 | HLB444 | 2.571 | 0.042 | HLB444 | 2.717 | 0.019 | Chow | -3.121 | 0.003 | Chow | -3.116 | 0.003 | Chow | -2.653 | 0.003 | - | - | - |
| **ko00510** | Glycan Biosynthesis and Metabolism | N-Glycan biosynthesis | - | - | - | - | - | - | - | - | - | - | - | - | Chow | -2.063 | 0.003 | Chow | -2.059 | 0.003 | Chow | -2.035 | 0.002 | Chow | -2.076 | 0.002 |
| **ko00601** | Glycan Biosynthesis and Metabolism | Glycosphingolipid biosynthesis - lacto and neolacto series | - | - | - | - | - | - | - | - | - | - | - | - | - | - | - | Chow | -2.282 | 0.003 | - | - | - | - | - | - |
| **ko00600** | Lipid Metabolism | Sphingolipid metabolism | B6 | -2.579 | 0.001 | B6 | -2.677 | 0.001 | - | - | - | - | - | - | - | - | - | - | - | - | HF | 2.443 | 0.002 | HF | 2.567 | 0.002 |
| **ko00061** | Lipid Metabolism | Fatty acid biosynthesis | B6 | -2.168 | 0.023 | B6 | -2.141 | 0.03 | - | - | - | - | - | - | Chow | -2.546 | 0.003 | Chow | -2.537 | 0.003 | Chow | -2.245 | 0.005 | Chow | -2.208 | 0.007 |
| **ko00564** | Lipid Metabolism | Glycerophospholipid metabolism | HLB444 | 2.344 | 0.001 | HLB444 | 2.161 | 0.003 | - | - | - | - | - | - | - | - | - | - | - | - | Chow | -2.201 | 0.003 | Chow | -2.065 | 0.007 |
| **ko00561** | Lipid Metabolism | Glycerolipid metabolism | - | - | - | - | - | - | - | - | - | B6 | -2.201 | 0.042 | HF | 2.58 | 0.003 | HF | 2.577 | 0.003 | HF | 2.381 | 0.003 | HF | 2.287 | 0.005 |
| **ko00072** | Lipid Metabolism | Synthesis and degradation of ketone bodies | - | - | - | - | - | - | - | - | - | - | - | - | HF | 2.002 | 0.003 | HF | 2.028 | 0.003 | - | - | - | - | - | - |
| **ko01040** | Lipid Metabolism | Biosynthesis of unsaturated fatty acids | - | - | - | - | - | - | - | - | - | - | - | - | - | - | - | - | - | - | Chow | -2.033 | 0.002 | - | - | - |
| **ko00120** | Lipid Metabolism | Primary bile acid biosynthesis | - | - | - | - | - | - | - | - | - | - | - | - | - | - | - | - | - | - | - | - | - | Chow | -2.094 | 0.002 |
| **ko00121** | Lipid Metabolism | Secondary bile acid biosynthesis | - | - | - | - | - | - | - | - | - | - | - | - | - | - | - | - | - | - | - | - | - | Chow | -2.094 | 0.002 |
| **ko02060** | Membrane Transport | Phosphotransferase system (PTS) | HLB444 | 2.627 | 0.013 | HLB444 | 2.608 | 0.03 | - | - | - | B6 | -2.249 | 0.042 | HF | 2.915 | 0.003 | HF | 2.921 | 0.003 | HF | 2.427 | 0.01 | HF | 2.396 | 0.007 |
| **ko02010** | Membrane Transport | ABC transporters | HLB444 | 3.413 | 0.001 | HLB444 | 3.433 | 0.001 | B6 | -3.098 | 0.042 | B6 | -3.255 | 0.028 | HF | 3.698 | 0.003 | HF | 3.689 | 0.003 | HF | 3.119 | 0.02 | - | - | - |
| **ko03070** | Membrane Transport | Bacterial secretion system | - | - | - | B6 | -2.023 | 0.03 | - | - | - | - | - | - | - | - | - | - | - | - | HF | 2 | 0.02 | HF | 2.134 | 0.007 |
| **ko00740** | Metabolism of Cofactors and Vitamins | Riboflavin metabolism | B6 | -2.273 | 0.007 | B6 | -2.062 | 0.039 | - | - | - | HLB444 | 2.351 | 0.028 | Chow | -2.635 | 0.003 | Chow | -2.611 | 0.003 | - | - | - | - | - | - |
| **ko00130** | Metabolism of Cofactors and Vitamins | Ubiquinone and other terpenoid-quinone biosynthesis | B6 | -2.461 | 0.002 | B6 | -2.544 | 0.002 | HLB444 | 2.228 | 0.042 | HLB444 | 2.385 | 0.019 | Chow | -2.738 | 0.003 | Chow | -2.724 | 0.003 | Chow | -2.081 | 0.037 | - | - | - |
| **ko00730** | Metabolism of Cofactors and Vitamins | Thiamine metabolism | B6 | -2.427 | 0.002 | B6 | -2.416 | 0.001 | HLB444 | 2.06 | 0.042 | HLB444 | 2.157 | 0.028 | Chow | -2.553 | 0.004 | Chow | -2.545 | 0.003 | - | - | - | - | - | - |
| **ko00790** | Metabolism of Cofactors and Vitamins | Folate biosynthesis | B6 | -2.563 | 0.002 | B6 | -2.6 | 0.002 | HLB444 | 2.497 | 0.019 | HLB444 | 2.581 | 0.012 | Chow | -2.917 | 0.003 | Chow | -2.911 | 0.003 | - | - | - | - | - | - |
| **ko00780** | Metabolism of Cofactors and Vitamins | Biotin metabolism | B6 | -2.416 | 0.003 | B6 | -2.473 | 0.002 | HLB444 | 2.297 | 0.028 | HLB444 | 2.441 | 0.019 | Chow | -2.773 | 0.003 | Chow | -2.77 | 0.003 | - | - | - | - | - | - |
| **ko00785** | Metabolism of Cofactors and Vitamins | Lipoic acid metabolism | B6 | -2.001 | 0.001 | B6 | -2.102 | 0.001 | - | - | - | - | - | - | - | - | - | - | - | - | - | - | - | HF | 2.101 | 0.002 |
| **ko00860** | Metabolism of Cofactors and Vitamins | Porphyrin and chlorophyll metabolism | B6 | -2.583 | 0.017 | B6 | -2.584 | 0.009 | B6 | -2.52 | 0.004 | B6 | -2.313 | 0.028 | HF | 2.931 | 0.003 | HF | 2.859 | 0.003 | HF | 2.976 | 0.002 | HF | 2.975 | 0.002 |
| **ko00770** | Metabolism of Cofactors and Vitamins | Pantothenate and CoA biosynthesis | B6 | -2.141 | 0.001 | B6 | -2.19 | 0.001 | - | - | - | - | - | - | - | - | - | - | - | - | - | - | - | HF | 2.012 | 0.002 |
| **ko00760** | Metabolism of Cofactors and Vitamins | Nicotinate and nicotinamide metabolism | - | - | - | - | - | - | HLB444 | 2.314 | 0.004 | HLB444 | 2.243 | 0.007 | Chow | -2.602 | 0.003 | Chow | -2.577 | 0.003 | Chow | -2.207 | 0.002 | Chow | -2.406 | 0.002 |

| | | | | | | | | | | | | | | | | | | | | | | | | | | |
|---|---|---|---|---|---|---|---|---|---|---|---|---|---|---|---|---|---|---|---|---|---|---|---|---|---|---|
| **ko00750** | Metabolism of Cofactors and Vitamins | Vitamin B6 metabolism | - | - | - | - | - | - | - | - | - | HLB444 | 2.055 | 0.012 | Chow | -2.348 | 0.003 | Chow | -2.337 | 0.003 | Chow | -2.088 | 0.002 | - | - | - |
| **ko00830** | Metabolism of Cofactors and Vitamins | Retinol metabolism | - | - | - | - | - | - | - | - | - | - | - | - | HF | 2.056 | 0.003 | HF | 2.07 | 0.003 | - | - | - | - | - | - |
| **ko00909** | Metabolism of Terpenoids and Polyketides | Sesquiterpenoid and triterpenoid biosynthesis | B6 | -2.124 | 0.001 | B6 | -2.342 | 0.001 | - | - | - | - | - | - | - | - | - | - | - | - | HF | 2.195 | 0.002 | HF | 2.398 | 0.002 |
| **ko00523** | Metabolism of Terpenoids and Polyketides | Polyketide sugar unit biosynthesis | - | - | - | - | - | - | - | - | - | HLB444 | 2.155 | 0.028 | Chow | -2.692 | 0.003 | Chow | -2.678 | 0.003 | Chow | -2.54 | 0.002 | Chow | -2.517 | 0.002 |
| **ko00908** | Metabolism of Terpenoids and Polyketides | Zeatin biosynthesis | - | - | - | - | - | - | - | - | - | - | - | - | Chow | -2.06 | 0.003 | Chow | -2.049 | 0.003 | - | - | - | - | - | - |
| **ko00900** | Metabolism of Terpenoids and Polyketides | Terpenoid backbone biosynthesis | - | - | - | - | - | - | - | - | - | - | - | - | Chow | -2.181 | 0.007 | Chow | -2.159 | 0.01 | Chow | -2.034 | 0.002 | Chow | -2.172 | 0.002 |
| **ko01051** | Metabolism of Terpenoids and Polyketides | Biosynthesis of ansamycins | - | - | - | - | - | - | - | - | - | - | - | - | HF | 2.258 | 0.003 | HF | 2.254 | 0.003 | - | - | - | - | - | - |
| **ko00903** | Metabolism of Terpenoids and Polyketides | Limonene and pinene degradation | - | - | - | - | - | - | - | - | - | - | - | - | HF | 2.007 | 0.003 | HF | 2.009 | 0.003 | - | - | - | - | - | - |
| **ko00230** | Nucleotide Metabolism | Purine metabolism | - | - | - | - | - | - | HLB444 | 2.752 | 0.004 | HLB444 | 2.652 | 0.007 | Chow | -2.901 | 0.003 | Chow | -2.849 | 0.003 | - | - | - | Chow | -2.414 | 0.037 |
| **ko00240** | Nucleotide Metabolism | Pyrimidine metabolism | - | - | - | - | - | - | HLB444 | 2.736 | 0.028 | HLB444 | 2.711 | 0.019 | Chow | -3.083 | 0.003 | Chow | -3.047 | 0.003 | Chow | -2.784 | 0.002 | Chow | -2.884 | 0.002 |
| **ko03410** | Replication and Repair | Base excision repair | HLB444 | 2.204 | 0.002 | HLB444 | 2.131 | 0.003 | - | - | - | - | - | - | Chow | -2.2 | 0.004 | Chow | -2.172 | 0.004 | Chow | -2.495 | 0.002 | Chow | -2.478 | 0.002 |
| **ko03030** | Replication and Repair | DNA replication | - | - | - | - | - | - | HLB444 | 2.179 | 0.019 | - | - | - | Chow | -2.424 | 0.007 | Chow | -2.41 | 0.003 | Chow | -2.124 | 0.01 | Chow | -2.307 | 0.002 |
| **ko03430** | Replication and Repair | Mismatch repair | - | - | - | - | - | - | HLB444 | 2.378 | 0.004 | HLB444 | 2.199 | 0.042 | Chow | -2.653 | 0.003 | Chow | -2.629 | 0.003 | Chow | -2.512 | 0.002 | Chow | -2.617 | 0.002 |
| **ko03440** | Replication and Repair | Homologous recombination | - | - | - | - | - | - | - | - | - | - | - | - | Chow | -2.279 | 0.015 | Chow | -2.284 | 0.015 | Chow | -2.149 | 0.005 | Chow | -2.412 | 0.002 |
| **ko03420** | Replication and Repair | Nucleotide excision repair | - | - | - | - | - | - | - | - | - | - | - | - | Chow | -2.189 | 0.003 | Chow | -2.214 | 0.003 | - | - | - | - | - | - |
| **ko02020** | Signal Transduction | Two-component system | HLB444 | 3.071 | 0.007 | HLB444 | 3.056 | 0.007 | B6 | -2.989 | 0.028 | B6 | -3.069 | 0.028 | HF | 3.475 | 0.004 | HF | 3.456 | 0.004 | HF | 2.974 | 0.027 | - | - | - |
| **ko03010** | Translation | Ribosome | B6 | -2.486 | 0.005 | B6 | -2.535 | 0.009 | HLB444 | 2.758 | 0.042 | - | - | - | Chow | -3.099 | 0.003 | Chow | -3.108 | 0.003 | Chow | -2.634 | 0.003 | Chow | -2.844 | 0.003 |
| **ko03013** | Translation | RNA transport | - | - | - | - | - | - | - | - | - | - | - | - | Chow | -2.32 | 0.003 | Chow | -2.294 | 0.003 | Chow | -2.202 | 0.002 | Chow | -2.218 | 0.002 |
| **ko03015** | Translation | mRNA surveillance pathway | - | - | - | - | - | - | - | - | - | - | - | - | Chow | -2.165 | 0.004 | Chow | -2.947 | 0.003 | - | - | - | - | - | - |
| **ko03008** | Translation | Ribosome biogenesis in eukaryotes | - | - | - | - | - | - | - | - | - | - | - | - | - | - | - | - | - | - | - | - | - | HF | 2.014 | 0.002 |
| **ko00362** | Xenobiotics Biodegredation and Metabolism | Benzoate degradation | - | - | - | - | - | - | B6 | -2.319 | 0.004 | B6 | -2.287 | 0.004 | HF | 2.466 | 0.003 | HF | 2.456 | 0.003 | - | - | - | - | - | - |
| **ko00791** | Xenobiotics Biodegredation and Metabolism | Atrazine degradation | - | - | - | - | - | - | HLB444 | 2.334 | 0.004 | HLB444 | 2.324 | 0.004 | Chow | -2.454 | 0.003 | Chow | -2.438 | 0.003 | - | - | - | - | - | - |
| **ko00983** | Xenobiotics Biodegredation and Metabolism | Drug metabolism - other enzymes | - | - | - | - | - | - | - | - | - | - | - | - | Chow | -2.601 | 0.003 | Chow | -2.572 | 0.003 | Chow | -2.626 | 0.002 | Chow | -2.672 | 0.002 |
| **ko00626** | Xenobiotics Biodegredation and Metabolism | Naphthalene degradation | - | - | - | - | - | - | - | - | - | - | - | - | HF | 2.062 | 0.003 | HF | 2.064 | 0.003 | - | - | - | - | - | - |
| **ko00363** | Xenobiotics Biodegredation and Metabolism | Bisphenol degradation | - | - | - | - | - | - | - | - | - | - | - | - | HF | 2.269 | 0.003 | HF | 2.292 | 0.015 | - | - | - | - | - | - |

| | | | | | | | | | | | | | | | | | | | | | | | | | | |
|---|---|---|---|---|---|---|---|---|---|---|---|---|---|---|---|---|---|---|---|---|---|---|---|---|---|---|
| **ko00633** | Xenobiotics Biodegredation and Metabolism | Nitrotoluene degradation | - | - | - | - | - | - | - | - | - | - | - | - | HF | 2.13 | 0.003 | HF | 2.127 | 0.003 | HF | 2.129 | 0.002 | HF | 2.151 | 0.002 |
| **ko00625** | Xenobiotics Biodegredation and Metabolism | Chloroalkane and chloroalkene degradation | - | - | - | - | - | - | - | - | - | - | - | - | - | - | - | HF | 2.009 | 0.003 | - | - | - | - | - | - |
